# Large-scale cellular-resolution read/write of activity enables discovery of cell types defined by complex circuit properties

**DOI:** 10.1101/2025.10.21.683734

**Authors:** Antonia Drinnenberg, Alexander Attinger, Allan Raventos, Charu Ramakrishnan, Barsin Eshaghi Gharagoz, Arjay Cordero, YoungJu Jo, La’Akea Siverts, Tanya L. Daigle, Bosiljka Tasic, Hongkui Zeng, Lisa M. Giocomo, Sean Quirin, Surya Ganguli, Karl Deisseroth

## Abstract

The complexity of the mammalian brain’s vast population of interconnected neurons poses a formidable challenge to elucidate its underlying mechanisms of coordination and computation. A key step forward will be technologies that can perform large-scale, cellular-resolution monitoring and interrogation of distributed brain circuit activity in behaving animals. Here, we present an all-optical strategy for precise optogenetic activity control of ∼10^3^ neurons and simultaneous activity monitoring of ∼10^4^ neurons within and across areas of mouse cortex—an order-of-magnitude leap beyond previous capabilities. Tracking population responses following delivery of precisely-defined widely-distributed activity patterns to the visual cortex of awake mice, we were surprised to identify neurons robustly responsive to stimulation of diverse ensembles, defying conventional like-to-like wiring rules. These cells were primarily deep L2/3 somatostatin-positive (SST) interneurons with functional properties distinct from other SST neurons, and appeared to play a role in brain dynamics that could only have been identified through broad cellular-resolution circuit interrogation. Our work reveals the value of measuring large-scale circuit-dynamical properties of functionally-resolved single cells, beyond genetic and anatomical classification, to define and explore the roles of cell types in brain function.

## INTRODUCTION

A major goal of modern neuroscience is to elucidate neural coding principles by discovering cause-and-effect relationships between neuronal activity and brain function. Two-photon (2P) all-optical technology has enabled precise control over the activity of ensembles of individual neurons in the intact mammalian brain, while simultaneously allowing readout of network dynamics and behavior^1,2^. Evolving from initial demonstrations of single-cell optogenetic 2P stimulation *in vivo*^3^ to including simultaneous Ca^2+^ activity readout^4^, this approach rapidly scaled towards the activation of tens of cells^5^—sufficient for influencing behavior in specific contexts^6–9^. By increasing the tissue volume accessible via optical means, continued efforts expanded the number of cells accessible for optical read-write control to hundreds within an acquisition volume^10–15^. With larger-scale optical access, greater numbers of neurons could be viewed to identify downstream stimulation-evoked activity changes in the surrounding, non-stimulated cells. This general approach has provided mechanistic insights into the functional impact of individual neurons^16^ or groups of individually-specified neurons^13,17–22^ on local network operation.

Despite these advances, major technological constraints have held back discovery. For example, current all-optical preparations typically allow readout from only a few hundred individually-specified neurons, and the observed signal strength is often compromised by conflicting requirements of expression level for the readout protein (typically a GCaMP-series Ca^2+^ indicator) versus the actuator protein (typically a light-gated membrane ion channel called a channelrhodopsin (ChR) encoded by an opsin gene). The readout indicator must be expressed at sufficiently high levels in order to achieve adequate signal strength, while the ChR must be expressed at relatively low levels in order to not excessively further increase protein expression burden imposed upon the cell by GCaMP expression, and to minimize incidental actuation from the GCaMP imaging light.

To date, intracranial adeno-associated virus (AAV) injection represents the most widely-used approach to drive opsin expression in all-optical preparations^23,24^, but such viral methods suffer from spatial inhomogeneity, expression changes over time, and high variability of transduction from cell to cell and across animals. Achieving stable and widespread presence of indicator and ChR with finely balanced expression levels has proven challenging^23,24^. Transgenic mouse line expression of both components could help address some of these challenges, but the single relevant transgenic line available (*Ai203*^25^)— expressing a soma-enriched ChR (ChroME) fused to GCaMP7s—requires higher stimulation power compared to preparations with more potent opsins^14^, which in turn constrains the number of neurons that can be stimulated simultaneously without eliciting thermal side effects^24,26^. Furthermore, while the membrane-bound indicator expression strategy dictated by this fusion protein strategy can be useful for specific applications (i.e. imaging compartmentalized Ca^2+^ signals in neurites), this design is less suited for population-level cellular activity imaging compared to cytosolic expression strategies^27^.

Here, we developed and employed a distinct all-optical read-write approach to address these challenges and advance neuroscience discovery; we targeted critical experimental barriers by jointly focusing on robust long-term all-optical access across cortex, reliable stimulation with minimal power, and activity readout technology with high signal-to-noise ratio (SNR). As part of this solution, we developed fully-transgenic all-optical tools and strategies yielding stable, well-balanced, and wide-spread co-expression of soma-enriched ChR and indicator, offering robust layer-specific read-write access across cortex without viral injection. Complementing this fully-transgenic approach to stimulating and recording from well-defined ensembles, we also developed a distinct “follower-focused” strategy that decouples transgenic opsin expression from dense viral indicator expression, to capture stimulation-evoked follower effects in the surrounding network, allowing read-write access to >1,000 neurons and read access to 10,000 neurons in a single acquisition volume and enabling the detection of hundreds of significantly modulated “follower” neurons per ensemble in the non-targeted population.

We took advantage of this rich view of evoked dynamics to gain insights into the single-cell resolution functional architecture of large-scale cortical networks, identifying a somatostatin-positive (SST) cell subtype in deep L2/3 with remarkable circuit-level recruitment properties. These cells, only identifiable using the new follower-focused approach, were termed general ensemble-response (GER) neurons because of their strikingly robust co-activation in response to a wide variety of unique large-scale stimulation patterns—a recruitment pattern that sharply contrasted with that of neighboring parvalbumin-positive (PV) or other SST interneuron subtypes. The broad response profile of GER neurons, discovered through large-scale optogenetic stimulation, extended to their functional properties during sensory stimulation and spontaneous activity: GER neurons in visual cortex were broadly responsive to a diverse range of visual stimuli and exhibited strongly synchronized firing during both visually evoked and spontaneous activity, suggesting they form a functional subnetwork. Modeling indicated that SST-GER neurons can achieve their broad and robust recruitment through a combination of high excitability and long-range spatial integration of excitatory inputs. This positions them to broadly monitor excitatory activity and, through the inhibitory action of SST neurons on the distal dendrites of pyramidal cells, gate the flow of top-down cortical information. Together, these discoveries and technologies help illuminate fundamental principles of cortical organization, revealing distinct roles for specific interneuron subtypes in shaping network responses and underscoring the value of broad circuit-level approaches for identification of functional cellular components of neural systems not discoverable by anatomy or gene expression alone.

## RESULTS

### Fully-transgenic all-optical strategies with layer-specificity and high sensitivity

We first generated two new transgenic reporter lines, designated here *Ai228* and *Ai229*, that express a soma-enriched form of the highly-sensitive excitatory channelrhodopsin ChRmine^13,28^ alongside GCaMP6m in a Cre-dependent manner (**Figure 1A**). *Ai228* (TIT2L-GCaMP6m-ICL1-ChRmine-TS-oScarlet-Kv2.1-ER-ICL2-IRES2-tTA2) is a TIGRE3.0 reporter^29^ whose construct contains: (1) a TRE-driven GCaMP6m cassette; (2) a CAG-driven ChRmine-oScarlet cassette, including a Kv2.1 motif^29^ for somatic enrichment of ChRmine-oScarlet expression; and (3) a CAG-driven tTA2 cassette for intrinsic activation of the TRE cassette^29^, including an IRES sequence positioned upstream of the tTA2 sequence to mitigate tTA2 expression. *Ai229* (TIT2L-GCaMP6m-IRES2-ChRmine-TS-Kv2.1-HA-ISL-tTA2) is a TIGRE2.0 reporter whose construct contains: (1) a TRE cassette with a bicistronic sequence to express GCaMP6m and soma-enriched ChRmine; and (2) an human synapsin promoter (hSyn, neuron-specific) driven tTA2 cassette. By excluding a red fluorophore in *Ai229*, we sought to facilitate separable cell-type or projection-specific labeling using the free red channel.

**Figure 1.**
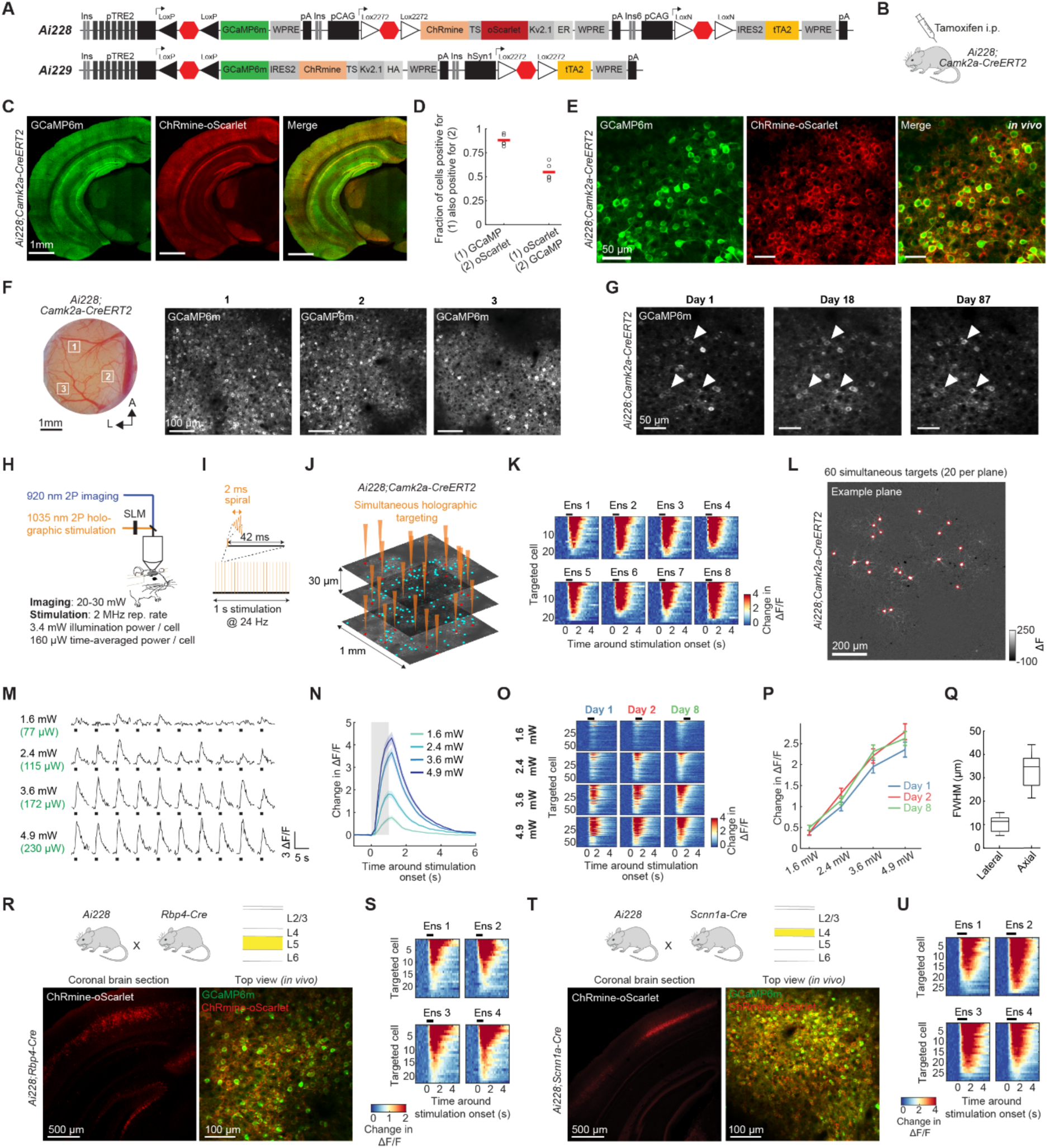
Fully-Transgenic All-Optical Strategies Employing the Potent Opsin ChRmine for Robust and Layer-Specific Read-Write Access Across Cortex. **A**, Schematic diagrams of the new transgenic reporters *Ai228* and *Ai229*. **B**, Double heterozygous, tamoxifen-induced *Ai228*;*Camk2a-CreERT2* mice were used for fully-transgenic all-optical experiments. **C**, Confocal images of a coronal brain section from a tamoxifen-induced *Ai228*;*Camk2a-CreERT2* mouse. Neurons throughout the forebrain express GCaMP6m (green) and soma-enriched ChRmine-oScarlet (red). **D**, Quantification of GCaMP6m and ChRmine-oScarlet co-expression in visual cortex of tamoxifen-induced *Ai228*;*Camk2a-CreERT2* mice (n = 5 FOVs, 2 mice). Red bars indicate mean values. **E**, Representative *in vivo* two-photon (2P) images acquired in layer 2/3 (L2/3, ∼150 µm from surface) posterior cortex of a tamoxifen-induced *Ai228*;*Camk2a-CreERT2* mouse. The distinct channels show expression of GCaMP6m (green) and soma-enriched ChRmine-oScarlet (red). **F**, Left: Photograph of a chronic cranial window over the dorsal posterior cortex of a tamoxifen-induced *Ai228*;*Camk2a-CreERT2* mouse. White rectangles outline the three fields-of-view (FOVs) shown on the right. A: anterior, L: lateral. Right: 2P images showing GCaMP6m expression in L2/3. **G**, 2P images of the same FOV in L2/3 visual cortex across time (Day 1, 18, and 87). Day 1 was acquired one week post-surgery. Arrowheads point to example neurons identified across timepoints. **H**, Schematic of all-optical experiments: simultaneous 2P Ca²⁺ imaging and 2P holographic stimulation were performed in the visual cortex of awake, head-fixed mice. SLM: spatial light modulator. Values represent the typical experimental settings used in *Ai228*;*Camk2a-CreERT2* mice. **I**, Schematic illustrating the temporal structure of a single stimulation trial: A user-defined 3D illumination pattern (“hologram”) consisting of single points generated by a SLM was scanned on the targeted neurons using a pair of galvo mirrors for 2 ms (spiral scan with 5 µm diameter, 5 revolutions). This low duty cycle of the stimulation minimizes data loss due to stimulation-light induced saturation of the image detector (“stimulation artifact”, **Figure S2A**). The stimulation was repeated at a chosen frequency (typically 24 and 32 Hz) during a stimulation period of 1 s or 1.5 s. **J**, 2P images of a 3-plane acquisition volume in L2/3 visual cortex from an *Ai228*;*Camk2a-CreERT2* mouse. GCaMP+ neurons distributed over the entire volume (cyan, n=192) were grouped into 8 ensembles of 24 simultaneously targeted neurons (data shown in **K**). Neurons of an example ensemble (“Ens 1”) are marked red. Orange triangles represent holographic stimulation. **K**, Heatmaps show the stimulation-evoked, trial-averaged change in activity in the targeted neurons (n = 5 trials per ensemble). Neurons are sorted by response amplitude. Black rectangle indicates the 1 s duration of the stimulation. Data from targets without an assigned ROI are not shown (**STAR Methods**). **L**, Trial-averaged ΔF image of an example plane during 1s stimulation of 60 simultaneous targets using 3.6 mW illumination power/neuron (10 trials, 32 Hz). Red circles indicate the 20 targeted neurons in this example plane. **M**, Single-trial activity in a representative neuron in response to varying stimulation powers (10 trials per power level, one power level per row, 32 Hz). Values on the left indicate illumination power/neuron (black) and time-averaged power/neuron (green, in brackets). Black rectangles: 1s stimulation period. **N**, Trial-averaged change in activity in response to varying stimulation powers averaged across targeted neurons (60 targets, 10 trials per power level, same ensemble as in **L**). Shaded area: SEM. **O**, Trial-averaged change in activity in targeted neurons in response to different power levels (rows) across days (columns). The same neurons were stimulated with the same hologram on Day 1, 2, and 8 (n = 60 simultaneous targets). Neurons are sorted by their average response across days for each power level (same sorting across days). **P**, Stimulation-evoked responses averaged across targeted neurons for different power levels (shown in **O**), split by recording day. Error bars indicate SEM. Using 3.6 mW, 53/60 targets were reliably activated on all three days. **Q**, Box plots of full-width half maximum (FWHM) values of the physiological point spread function (pPSF) for individual neurons in L2/3 of an *Ai228*;*Camk2a-CreERT2* mouse. Lateral FWHM (µm): range 5.3-15.1, median 11.4 (n = 10 trials, n = 37 neurons from 1 mouse). Axial FWHM (µm): range 22-46, median 34.9 (n = 10 trials, 28 neurons from 1 mouse). See also **Figures S2G and S2H**. Values are comparable to viral approaches expressing soma-enriched ChRmine^13^. **R**, Top: *Ai228* mice were crossed to *Rbp4-Cre* mice to target expression of GCaMP6m and soma-enriched ChRmine-oScarlet to layer 5 (L5) excitatory neurons. Bottom: Confocal image of a coronal section (left) and *in vivo* 2P image of L5 dorsal cortex (right) of an *Ai228*;*Rbp4-Cre* mouse. Neurons in L5 express GCaMP6m (green) and soma-enriched ChRmine-oScarlet (red). **S**, Heatmaps show the stimulation-evoked, trial-averaged change in activity in targeted L5 neurons of an *Ai228*;*Rbp4-Cre* mouse (4 ensembles, 24 targets each, 3-plane volume with 30 µm spacing and 1x1 mm^2^ FOV). Stimulation: 1s, 24 Hz, 5.8 mW illumination power/neuron, 278 µW time-averaged power/neuron, 10 trials/ensemble. **T**, As in **R**, but using the Cre driver *Scnn1a-Cre* to target expression to excitatory neurons in layer 4 (L4). **U**, As in **S**, but for targeted L4 neurons of an *Ai228*;*Scnn1a-Cre* mouse (4 ensembles, 30 targets each). Stimulation: 1.5 s, 32 Hz, 2.8 mW illumination power/neuron,179 µW time-averaged power/neuron, 25 trials/ensemble.

We next employed diverse Cre-delivery strategies to evaluate *Ai228* and *Ai229* (**Table S1**). For forebrain-wide co-expression of GCaMP6m and ChRmine without necessitating viral injection, we crossed *Ai228* mice with the inducible *Camk2a-CreERT2* Cre driver line^30^. Tamoxifen-induced *Camk2a-CreERT2;Ai228* mice expressed GCaMP6m and soma-enriched ChRmine-oScarlet throughout the forebrain (**Figures 1B-1C and S1A-S1C**), potentially providing all-optical access to excitatory neurons in the cortex and the hippocampus. Within the cortex, 88 ± 2.7% (mean ± SEM) of GCaMP+ neurons also expressed ChRmine-oScarlet, while 55 ± 4.1% of ChRmine-oScarlet+ neurons also expressed GCaMP (**Figure 1D**). The lower percentage of ChRmine-oScarlet+ neurons also expressing GCaMP is possibly due to TRE silencing, a reported effect in related TIGRE lines^25,29^. To quantify the percentage of labeled neurons in *Camk2a-CreERT2;Ai228*, we used the neuronal marker NeuN (**Figures S1D and S1E**). 41 ± 2.2% of NeuN+ neurons expressed ChRmine-oScarlet, while 26 ± 2.0% of NeuN+ neurons expressed GCaMP. As excitatory neurons represent approximately 80% of all neurons in cortex, we estimate that in tamoxifen-induced *Camk2a-CreERT2*;*Ai228* mice, approximately one-third (32.5%) of the excitatory cortical population express GCaMP6m.

We next implanted chronic cranial windows in *Camk2a-CreERT2;Ai228* mice for long-term all-optical access across the posterior dorsal cortex (**Figure 1E-F and S1F**). Preparations remained stable for months (**Figure 1G**), in stark contrast to virally expressed, soma-enriched ChRmine and GCaMP6m, which typically degrade within weeks post-injection (**Figure S1G**). To evaluate the all-optical performance of the *Camk2a-CreERT2;Ai228* approach, we targeted ensembles of excitatory neurons in L2/3 visual cortex for holographic photostimulation employing a 2P microscopy system equipped with a spatial light modulator (SLM) (**Figures 1H and 1I, STAR Methods**). Based on the fraction of GCaMP+ neurons expressing ChRmine-oScarlet (**Figure 1D**), approximately 88% of GCaMP+ neurons should be photo-activatable. Indeed, when manually selecting GCaMP+ neurons in a 3-plane volume (1x1 mm² field-of-view (FOV), 30 µm axial spacing) and stimulating ensembles each comprised of 24 simultaneously stimulated targets, we achieved an 86% response rate using minimal stimulation power (3.4 mW illumination power/neuron, 0.16 mW time-averaged power/neuron, 24 Hz stimulation, 2 ms light exposure per pulse, stimulation laser operating at 2 MHz repetition rate). Thus, in *Camk2a-CreERT2*;*Ai228*, we found that neurons can be effectively stimulated with low power (and conveniently, for a success rate close to 90%, target selection can be simplified by relying solely on the GCaMP6m signal).

We further evaluated this approach by stimulating 60 simultaneous targets at varying power and frequency (**Figures 1L-1N and S2B-S2F**). As expected, stimulation-evoked responses increased with power and frequency. The responses were remarkably stable across trials within a session (average response correlation: 0.76; n = 10 trials per condition, **Figures 1M and S2B**). When stimulating the same ensemble with the same hologram across days (Days 1, 2, and 8), the targeting efficiency remained high (**Figures 1O and 1P**), demonstrating suitability of this approach for longitudinal all-optical experiments. Furthermore, we found that the *Camk2a-CreERT2;Ai228* approach allowed for photo-targeting with high spatial resolution; the full-width half maximum (FWHM) of the physiological point-spread function (pPSF) was 11.4 µm laterally, and 34.9 µm axially (**Figures 1Q, S2G and S2H**) and when targeting two closely spaced ensembles, neurons responded exclusively when directly targeted (**Figures S2I and S2J**). Cross-stimulation (i.e. unintended activation of ChR by the imaging laser) and signal strength was markedly improved in *Camk2a-CreERT2*;*Ai228* mice compared to the bicistronic viral approach for expressing soma-enriched ChRmine and GCaMP6m employed in this reference^13^ (**Figure S3**). This improved performance is likely a consequence of lower ChRmine but higher GCaMP6m expression with the transgenic strategy.

For layer-specific fully-transgenic all-optical access, we crossed *Ai228* to the layer 4 (L4) Cre driver *Scnn1a-Cre*^30^ and the layer 5 (L5) Cre driver *Rbp4-Cre*^31^. Both approaches yielded robust co-expression of GCaMP6m and ChRmine-oScarlet in the targeted layers enabling reliable holographic stimulation of L4 and L5 ensembles with remarkably low power (illumination power/neuron: L4: 2.8 mW, L5: 5.8 mW; **Figures 1R-1U and S4A-F**). For applications that require simplified breeding schemes, we also explored viral Cre delivery in *Ai228* mice. While intracranial AAV injections failed to produce stable expression (**Table S1**), systemic delivery of a PHP.eB-coated AAV containing the pan-excitatory enhancer eHGT_78h^32^ and a PEST sequence to attenuate Cre expression achieved stable co-expression of GCaMP6m and soma-enriched ChRmine-oScarlet across cortex (**Figure S4G**).

To facilitate fully-transgenic all-optical experiments with the red channel available for additional labeling, we evaluated *Ai229* mice (**Table S1, Figure S5**). *Camk2a-CreERT2*;*Ai229* mice expressed GCaMP6m and soma-enriched, HA-tagged ChRmine in excitatory neurons across the forebrain (**Figures S5A-E**). Similar to *Ai228*, *Ai229* permitted reliable stimulation of GCaMP+ neurons with minimal power (**Figures S5F-I**), demonstrating utility for all-optical experiments. However, likely due to a higher ChRmine:GCaMP expression ratio resulting from the bicistronic design, *Ai229* exhibited slightly increased cross-stimulation and reduced SNR compared to *Ai228* (**Figure S3**).

### Order-of-magnitude scaling up of all-optical access

The fully-transgenic approaches described above provide robust and convenient read-write access across cortex. However, for a more complete view of the downstream activity changes in the surrounding network, monitoring the entire neuronal population within a given FOV would be advantageous. Achieving such dense readout via transgenic indicator expression is challenging because sufficient transgenic expression levels require transcriptional amplification of the tTA/TRE-system^33^, but the density of TRE-driven expression is limited by cell-type-specific TRE silencing^25,29^.

We therefore developed a novel “follower-focused” strategy, which combines transgenic opsin expression targeted to a specific cell class with dense, pan-neuronal indicator labeling. To implement this new approach, we first evaluated differences in transgenic ChRmine expression between two Cre-dependent ChRmine-oScarlet TIGRE2.0 reporter lines, *Ai219* and *Ai230* (**Figures S6A-S6G and Table S1**). *Ai219* is our newly developed reporter featuring TRE-driven, soma-enriched ChRmine-oScarlet expression, while in contrast, *Ai230* expresses soma-enriched ChRmine-oScarlet driven by the CAG promoter^34^. We observed very high expression levels per cell in *Ai219* (in line with the potent activity of the TRE promoter compared to CAG) and found that, across various Cre delivery strategies, the density of labeled cells was sparse compared to *Ai230*, possibly due to TRE silencing in *Ai219*. Furthermore, we observed that some crosses of *Ai219* with Cre drivers led to embryonic lethality, preventing the acquisition of double-positive offspring, or to abnormal brain morphology (details listed in **Table S1**). While *Ai219* might be useful for specific applications that benefit from higher per-cell expression, each Cre delivery strategy requires careful evaluation for potential abnormalities. For our experiments, we opted for the lower and more widespread expression achieved by CAG-driven ChRmine-oScarlet in *Ai230*.

We crossed *Ai230* mice with either the excitatory Cre driver *Slc17a7-Cre*^35^ or *Camk2a-CreERT2* mice (**Figures 2A, S6B and S6C**). Double heterozygous mice were intracranially injected with an AAV expressing hsyn-driven GCaMP8s, with GCaMP8s restricted to the soma via ribosome-tagging^36^, in an optimized construct (**STAR Methods**) to maximize expression level. The resulting brain preparations exhibited soma-enriched ChRmine-oScarlet expression in excitatory neurons and ribosome-tagged GCaMP8s (“riboGCaMP8s”) in both excitatory and inhibitory neurons (assessed with *in vivo* 2P imaging; **Figure 2B and 2C**). The dense indicator expression with reduced neuropil labeling allowed us to detect ∼2,000 neurons across a single 1x1 mm² FOV (**Figure 2D**). In a single acquisition volume (1000 x 1000 x 120 µm³, across 5 planes with 30 µm axial spacing), we demonstrated read-access to ∼10,000 neurons (9,978 GCaMP-ROIs) and read-write-access to >1,000 neurons using minimal stimulation power (successful stimulation of 1,053 GCaMP+ and oScarlet+ neurons using 2.7 mW illumination power/neuron, 173 µW time-averaged power/neuron, 60 targets/ensemble, **Figures 2E, 2F and S6H**).

**Figure 2.**
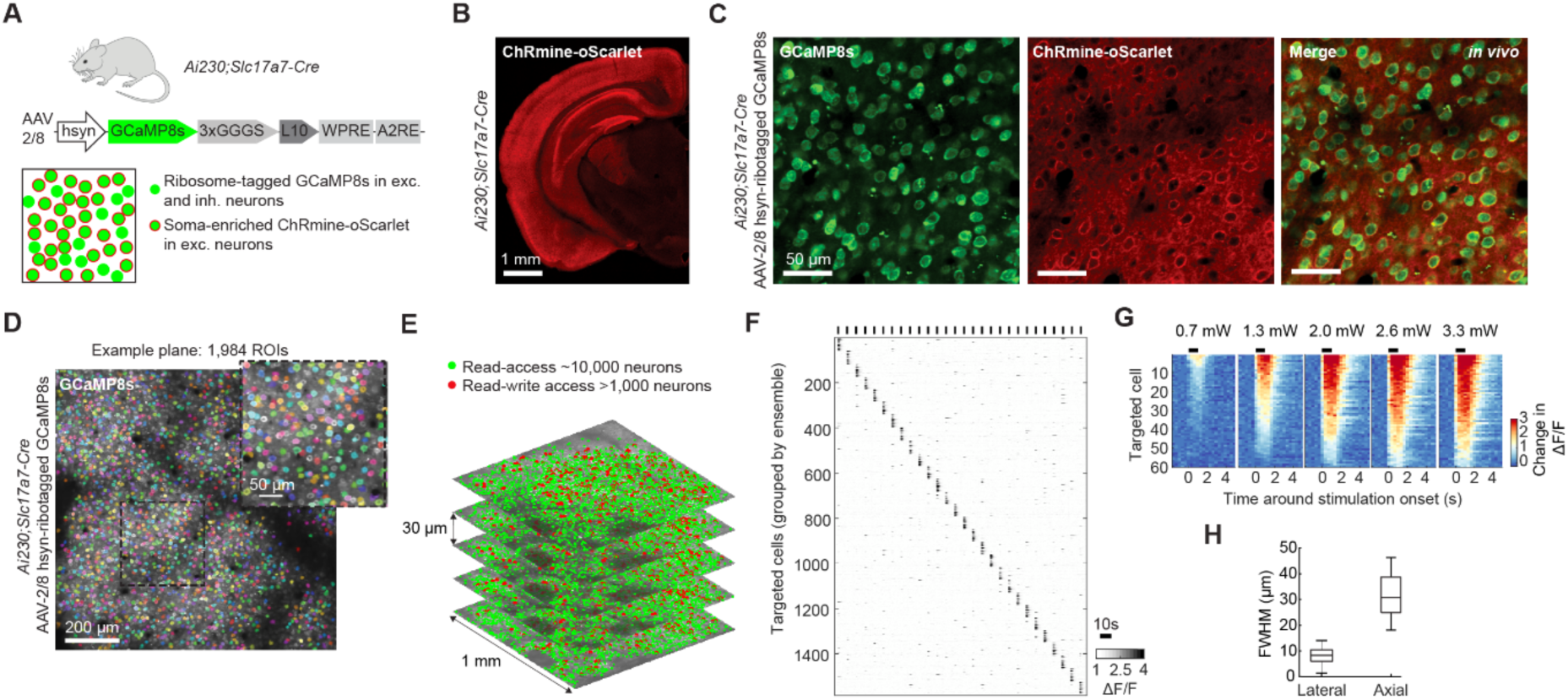
Scaled-Up All-Optical Experiments with Read/Write Access to >1,000 Neurons and Read Access to 10,000 Neurons. **A**, Schematic illustrating the “follower-focused” approach, which combines transgenic ChRmine expression in excitatory neurons with dense, virally-driven expression of ribosome-tagged GCaMP8s in excitatory and inhibitory neurons to provide a more complete view of downstream follower effects evoked by holographic stimulation. For this approach, *Ai230* mice (a Cre-dependent reporter for soma-enriched ChRmine-oScarlet^34^) were crossed with either *Slc17a7-Cre* mice or *Camk2a-CreERT2* mice (both excitatory Cre driver). Double heterozygous mice were intracranially injected with AAV for hsyn-driven ribosome-tagged GCaMP8s expression (pan-neuronal “riboGCaMP8s”), in an optimized construct (**STAR Methods**) to maximize expression level. **B**, Confocal image of a representative coronal brain section from an *Ai230*;*Slc17a7-Cre* mouse, showing soma-enriched ChRmine-oScarlet (red) across the forebrain. **C**, Representative *in vivo* 2P images of L2/3 visual cortex from an *Ai230*;*Slc17a7-Cre* mouse with pan-neuronal riboGCaMP8s, showing dense riboGCaMP8s labeling in excitatory and inhibitory neurons (green) and soma-enriched ChRmine-oScarlet expression targeted to excitatory neurons (red). **D**, Representative 2P image of L2/3 visual cortex from an *Ai230*;*Slc17a7-Cre* mouse with pan-neuronal riboGCaMP8s (gray), with Suite2p^76^-extracted regions-of-interest (ROIs) overlaid. Inset: Zoomed-in view of outlined area (307 µm x 307 µm). **E**, 2P images of a representative 5-plane imaging volume in an *Ai230*;*Slc17a7-Cre* mouse with pan-neuronal riboGCaMP8s. 9,978 GCaMP-ROIs (green) were imaged. 1,680 GCaMP-ROIs were also oScarlet+ (red) and subsequently targeted for stimulation (28 ensembles, 60 targets/ensemble, 2.7 mW illumination power/neuron, 173 µW time-averaged power/neuron, 32 Hz, responses shown in **F**). **F**, Stimulation-evoked responses of the targeted neurons shown in **E**. Heatmap shows the trial-averaged ΔF/F activity (n = 5 trials). Neurons are grouped by ensemble identity. 1,053 neurons were successfully stimulated. Data from targets without an assigned ROI are not shown (**STAR Methods**). Black bars on top indicate the start of the 1.5 s stimulation period for each of the 28 ensembles. **G**, Trial-averaged change in activity of a representative ensemble in response to varying stimulation powers. Targets (n = 60 neurons) were selected from the pool of responsive neurons of experiment shown in **E**. Stimulation: 1.5 s, 32 Hz, 10 trials. Values indicate illumination power/neuron. Percentages of successfully stimulated neurons (**STAR Methods**) for each power level: 26%, 80%, 90%, 92%, 92%. **H**, Box plot of FWHM values of the pPSF for individual neurons in L2/3 of an *Ai230*;*Slc17a7-Cre* mouse with pan-neuronal riboGCaMP8s. Lateral FWHM (µm): range 1.9-14.6, median 8.7 (n = 10 trials, n = 28 neurons from one mouse). Axial FWHM (µm): range 18-65, median 30.8 (n = 10 trials, n = 48 neurons from one mouse).

We further evaluated the *Ai230* ribo-GCaMP8s approach by examining power-response relationships (**Figures 2G and S6I**) and stimulating large ensembles (**Figure S6J**). Stimulation was achieved with high spatial precision (pPSF with a FWHM of 8.7 µm laterally and 30.8 µm axially, **Figures 2H, S2K and S2L**) and minimal cross-stimulation (**Figure S3**). Similar to the fully-transgenic approach, preparations remained remarkably stable across time, enabling tracking of individual neurons over many weeks (**Figure S6M**) and permitting all-optical experiments in individual animals for up to 70 weeks (**Figures S6N-S6R**). Thus, the new follower-focused approach (transgenic ChRmine combined with pan-neuronal riboGCaMP8s) allowed order-of-magnitude scaling-up of all-optical experiments in cortex, enabling long-term stimulation access to large cortical ensembles combined with dense sampling of thousands of neurons across large cortical volumes for flexible and versatile applications (for example, this approach is readily compatible with use of inter-areal cortical FOVs such as those spanning V1 and higher visual areas; **Figures S7A and S7B**).

### Widespread specific activity patterns evoked by holographic stimulation

We hypothesized that the improved readout conferred by the follower-focused approach—due to the minimized neuropil contamination, more sensitive indicator, increased number of imaged neurons, and access to both excitatory and inhibitory populations—could powerfully enable insight into the downstream activity patterns evoked by precise circuit manipulations. We tested this possibility by stimulating large ensembles of excitatory neurons in L2/3 of primary visual cortex (V1, identified by retinotopic mapping, **Figures S7A and S7B**) in awake mice. The ensembles consisted of a median of 50 targets per ensemble (n = 104 ensembles across 10 mice, **Figure S7C**). Target neurons were either randomly selected (“random”), or chosen based on specific visual-stimulus preference selectivity (“functional”). For the construction of functional ensembles, neurons were chosen based on imaging sessions performed prior to the stimulation experiments; the functional ensembles either included neurons tuned for a specific stimulus orientation (“orientation-selective”), or neurons responsive to a specific set of natural scenes (“scene-selective”; ensembles selected based on hierarchical clustering of scene-evoked responses). Ensembles were stimulated for 1.5 s at 32 Hz (**STAR Methods**) while mice were watching a gray screen. Across ensembles, 84.5 ± 0.95% (mean ± SEM) of the targeted neurons were successfully stimulated. Monitoring the stimulation-evoked activity of thousands of neurons per acquisition volume (median 8,318 ROIs, range 4,962-10,525), we observed widespread downstream effects in the non-targeted population, including far from stimulation targets (**Figures 3A, 3B and S7D**). The evoked changes in activity of these downstream neurons (including populations that were consistently excited and other populations that were consistently inhibited) were found to exhibit large ΔF/F amplitudes and trial-by-trial reliability (**Figure 3B**).

**Figure 3.**
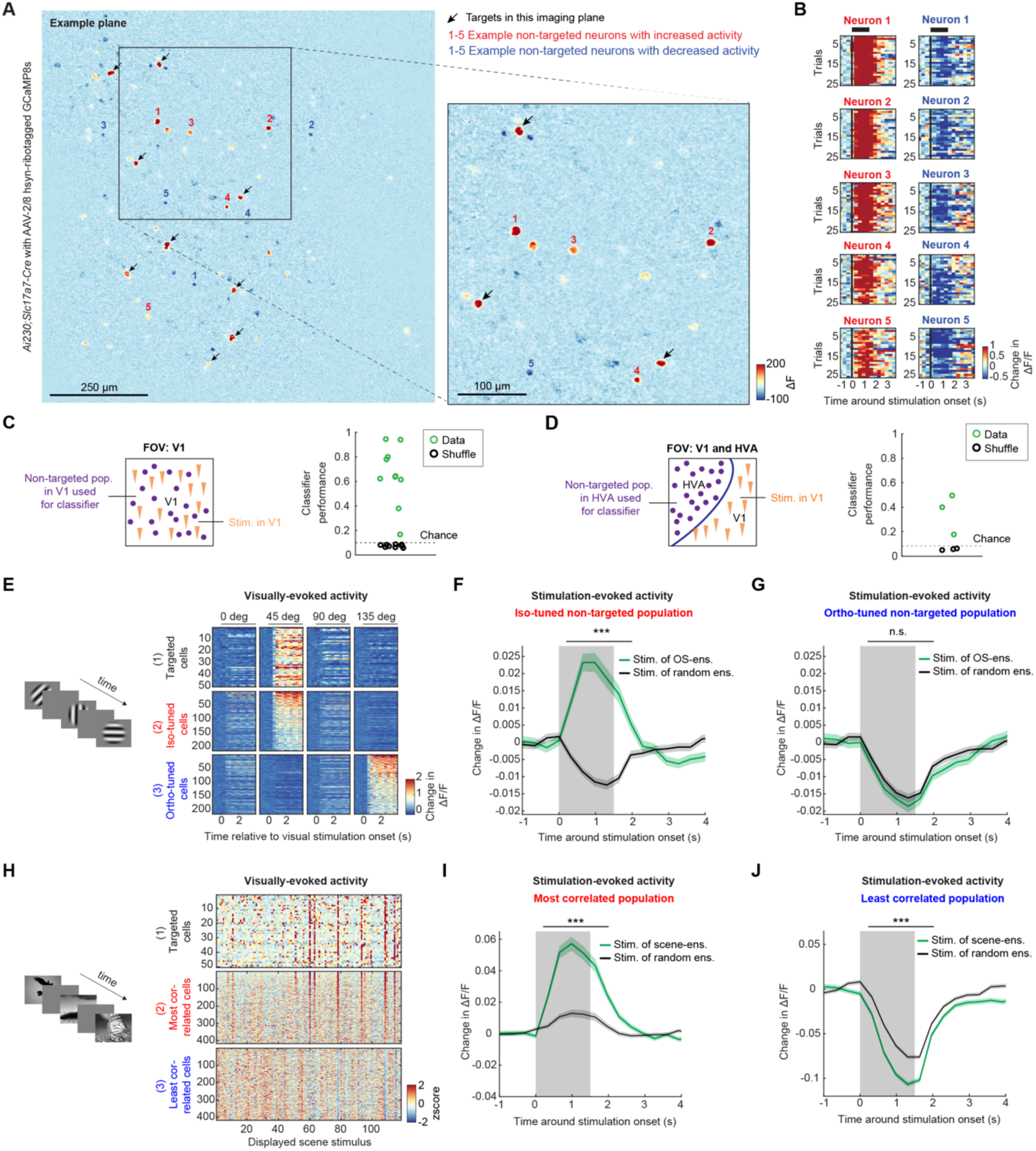
Widespread and Specific Downstream Activity Patterns Evoked by Large-Scale Holographic Stimulation Within and Across Areas of Visual Cortex. **A**, Trial-averaged ΔF images of an example plane during stimulation of 50 excitatory neurons in L2/3 visual cortex from an awake *Ai230*;*Slc17a7-Cre* mouse with pan-neuronal riboGCaMP8s (“follower-focused approach”). Stimulation: 1.5 s, 32 Hz, 25 trials. Left: Entire plane. Right: Zoom in. Arrows: Targets within this imaging plane (n = 9). 41 targets were located in the other four planes (locations shown in **Figure S7D**). Numbers correspond to the neurons shown in **B**. **B**, Single-trial change in activity of representative non-targeted neurons that increased (left) or decreased (right) their activity upon stimulation. **C**, Left: Schematic of classifier analysis. Stimulation was performed in primary visual cortex (V1, identified by retinotopic mapping, **Figure S7A**). A support-vector machine classifier was trained on the stimulation-evoked responses of the non-targeted V1 neurons (targets and nearby neurons excluded, **STAR Methods**). Right: Average classifier performance across ensembles for 10 mice (V1 dataset, 10-12 ensembles per mouse, **Figure S7C**). Each point represents the average performance for a single mouse. “Shuffle”: performance for shuffled trial labels. The dashed line indicates the maximum chance level across mice (0.1, based on 10-12 ensembles per mouse). See also **Figures S8A-C**. **D**, Same as **C** but for inter-areal experiments. The FOV was positioned to include both V1 and a higher visual area (HVA, identified by retinotopic mapping). Stimulated neurons were located in V1. The classifier was exclusively trained on stimulation-evoked responses of the non-targeted population in the HVA (3 mice, 12-13 ensembles per mouse, **Figure S8D**). The dashed line indicates the maximum chance level across mice (0.08, based on 12-13 ensembles per mouse). **E**, Left: Schematic of the visual grating stimuli. Each stimulus was shown for 4 s with 4 s inter-stimulus interval. Right: Trial-averaged ΔF/F responses for three groups of L2/3 neurons in V1 from an example mouse to the different gratings (0°, 45°, 90°, 135°). (1) Targeted cells: A representative orientation-selective (OS) stimulation ensemble (n = 50 cells). (2) Iso-tuned cells: the top 5% of neurons in the non-targeted population with most similar orientation preference to this stimulation ensemble. (3) Ortho-tuned cells: the top 5% of neurons with most orthogonal orientation preference. **F**, Average stimulation-evoked responses of non-targeted neurons with orientation tuning most similar to the stimulated ensembles (selected as the top 5% of iso-tuned neurons per ensemble) during the stimulation of OS-ensembles (orange) and size-matched random ensembles (control, black) (n = 6,150 neurons from 8 mice, shading indicates SEM). Responses to OS-ensembles were significantly higher compared to control (n = 31 OS-ensembles, control: average of 4 random ensembles per mouse, paired t-test, p = 2.9 x 10⁻⁴). Gray shading: stimulation. Black bar: time window used for quantification. **G**, Same as **F** but for the top 5% ortho-tuned neurons per ensemble. Responses evoked by OS-ensembles were not significantly different from responses to random ensembles (n = 31 OS-ensembles, control: average of 4 random ensembles per mouse, paired t-test, p = 0.64). **H**, Left: Schematic of the natural scene stimuli. Individual scenes were presented for 0.75 s with 1.75 s inter-stimulus interval. Right: Z-scored, trial-averaged responses to 118 scene stimuli in an example mouse are shown for: (1) a representative scene-selective stimulation ensemble (n = 50 cells), (2) the top 5% most similar non-targeted neurons, and (3) the top 5% least similar non-targeted neurons. Similarity was defined as the correlation between the trial-averaged response vector of each non-targeted neuron and the mean trial-averaged response vector of the entire stimulation ensemble. **I**, Average stimulation-evoked responses of the top 5% most similar neurons per ensemble during stimulation of scene-selective ensembles (orange) and size-matched random ensembles (control, black) (n = 3,706 neurons from 7 mice). Responses to scene-selective ensembles are significantly higher compared to control (n = 19 scene-selective ensembles, control: average of 4 random ensembles per mouse, paired t-test, p = 2.0 x 10^-3^). **J**, Same as **I** but for the top 5% least similar neurons in the non-targeted population. Responses evoked by scene-ensembles are significantly lower compared to those evoked by random ensembles (n = 19 scene-selective ensembles, control: average of 4 random ensembles per mouse, paired t-test, p = 5.0 x 10^-3^).

We assessed whether stimulation ensemble identity could be decoded from recruited activity in downstream neurons. We trained a support-vector machine classifier to decode ensemble identity using the single-trial, stimulation-evoked responses of the non-targeted neurons (defined as all neurons within the acquisition volume, excluding both targeted neurons and nearby neurons, **STAR Methods**), with 80% of the trials serving as training set and 20% as the test set (5-fold cross-validation). The classifier successfully predicted ensemble identity above chance (**Figures 3C and S8A**). Critically, the decoding performance was at chance level before stimulation onset (**Figure S8B**) and when shuffling trial labels (**Figure 3C**, black circles), indicating that the observed decoding performance was not artificially inflated by classifier overfitting. Intriguingly, decoder performance was as successful with stimulation ensembles that were randomly-defined compared with stimulation ensembles that were functionally-defined (**Figure S8C**). In an additional set of experiments (**Figure S8D**, n = 3 mice), we simultaneously recorded activity in V1 and in an adjacent higher visual area (HVA, identified by retinotopic mapping, **Figure S7A**) and investigated whether stimulation ensemble identity could also be decoded from evoked activity that propagates across cortical areas. We stimulated ensembles restricted to V1 and used exclusively the non-targeted population in the HVA to train and evaluate the classifier. Notably, the classifier performed above chance also in these inter-areal experiments (**Figures 3D and S8E**). Thus, stimulating large excitatory ensembles in L2/3 visual cortex elicits a reliably informative network-wide response in downstream non-targeted neurons, within and across visual areas. These changes are not random, but rather encode the identity of the stimulated ensemble, creating a unique footprint in the pattern of recruited neurons.

As V1 circuits exhibit “like-to-like” connectivity patterns, where neurons with similar visual properties are more likely to be connected^37–39^, we asked if the stimulation of functional ensembles with specific visual properties would selectively influence those non-targeted neurons that share these properties^13,20^. We found that stimulation of orientation-selective ensembles indeed specifically increased the activity of neurons in the non-targeted population with the most similar tuning (“iso-tuned” neurons), while neurons with orthogonal tuning were suppressed (**Figures 3E-3G**). In contrast, both groups of neurons were suppressed upon stimulation of size-matched, random ensembles. We note that the specific increase in activity in iso-tuned non-targeted neurons, but not ortho-tuned non-targeted neurons, upon stimulation of orientation-selective ensembles was maintained over remarkable distances (>150 µm) from the nearest stimulation target (**Figures S8F and S8G**). This greatly exceeds the spatial range of such orientation-specific recruitment in the non-targeted population (<30 µm) reported in earlier all-optical studies of V1^20^. Furthermore, we detected functionally-selective recruitment for scene-tuned ensembles: non-targeted neurons exhibiting the strongest correlation with the stimulated scene-tuned ensemble showed increased activity, while those with the lowest correlation decreased their activity (**Figures 3H-3J**). These results align with previous studies identifying highly correlated responses to natural scenes as a strong predictor of connection probability^37^.

### General ensemble-response properties in a subset of downstream neurons

This functionally selective recruitment of neurons, along with the successful decoding of ensemble identity from downstream activity of non-targeted neurons, indicate that the activity patterns in the non-targeted (follower) population evoked by different large-scale stimulated ensembles are distinct. However, we also surprisingly observed that a subset of non-targeted neurons was exhibiting more general response patterns, becoming activated by stimulation of any of several ensembles—even though targeted neurons of different ensembles were completely non-overlapping (**Figures 4A and 4B**). These general ensemble-response (GER) neurons were recruited reliably across trials, and their activation was observed even when the nearest stimulation target was >150 µm away (**Figure S9A**).

**Figure 4.**
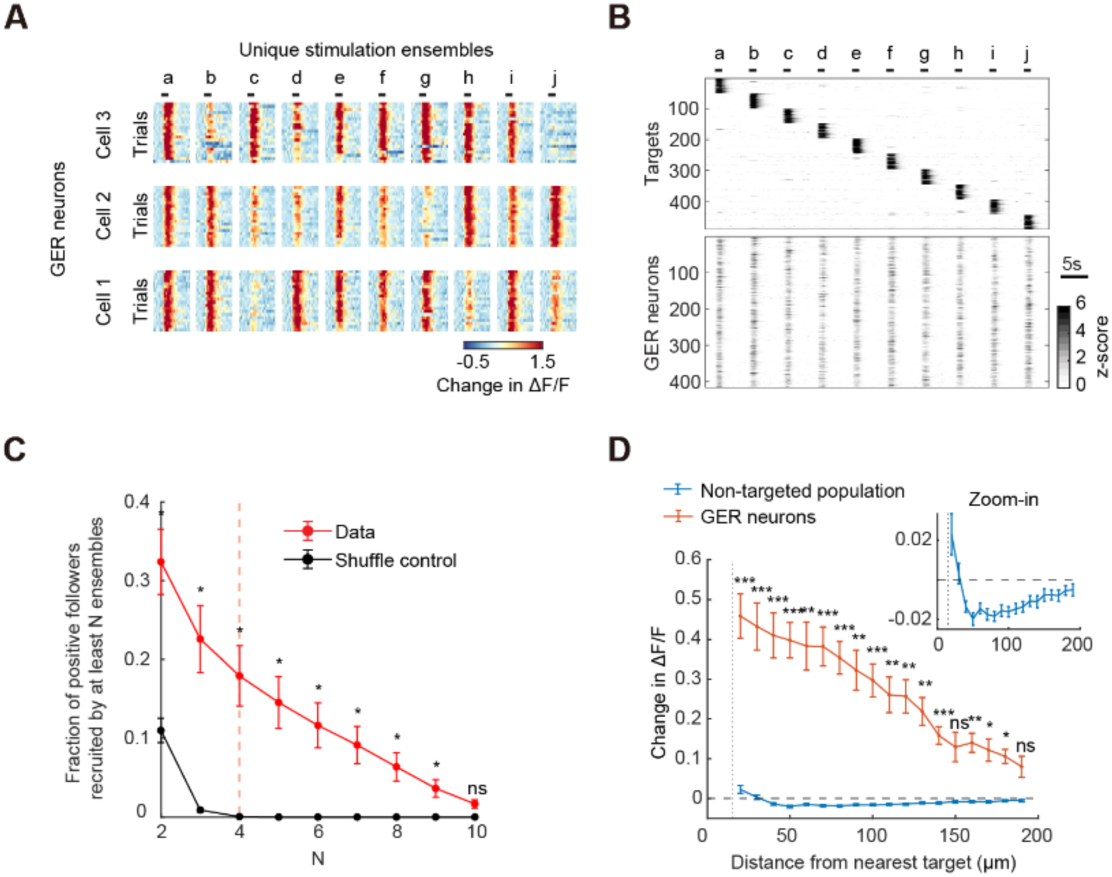
Unique and Robust General Recruitment Patterns Observed in a Subset of Follower Neurons. **A**, Single-trial change in activity of three example non-targeted neurons that were co-activated during stimulation by many of the 10 different, non-overlapping (disjoint) ensembles (ensembles a-j; 50 randomly-selected targets per ensemble, 32 Hz stimulation). Individual heatmaps show the reliable responses across trials (20 trials per ensemble), from -2 s before to 6 s after stimulation onset. Black bars indicate the 1.5 s stimulation period. Preparation: *Ai230*;*Slc17a7-Cre* mouse with pan-neuronal riboGCaMP8s. **B**, Trial-averaged stimulation-evoked responses recorded in the same example stimulation experiment as in **A**. Data are shown for the targeted neurons (top, sorted by ensemble identity) and for non-targeted neurons with significantly increased activity (Student’s t-test p <0.005) in response to stimulation of at least 4 different ensembles (bottom, n = 416 neurons). Hereafter this population of non-targeted neurons is referred to as general ensemble-response (GER) neurons. **C**, Fraction of positive followers (non-targeted neurons with significantly increased activity, Student’s t-test p <0.005) recruited by at least N ensembles as a function of N, for experimental data (V1 dataset, n = 10 mice, red) or in a shuffle control (black). Wilcoxon signed-rank tests (Bonferroni-corrected) assessed data vs. control (***p≤ 0.001, **p≤ 0.01, *p≤ 0.05, ns p>0.05). Dashed orange line: threshold for classifying neurons as GER neurons, defined as recruited by ≥4 ensembles. **D**, Stimulation-evoked change in activity as a function of distance to nearest target for GER neurons (orange) and all non-targeted neurons (blue), presented as mean ± SEM for n = 10 mice. For this analysis, GER neurons were defined on a left-out set of ensembles (**STAR Methods**). One-sample t-tests (Bonferroni-corrected) assessed responses of GER cells above zero. Horizontal dashed line: zero line. Inset: Zoom in on the data of all non-targeted neurons.

To systematically study this unexpected phenomenon, we first identified neurons within the non-targeted population with significant change (Student’s t-test p <0.005) in stimulation-evoked activity (using the V1 dataset, n =10 mice, 10-12 ensembles per mouse, **Figure S7C**). We found that stimulation of individual neuronal ensembles (median of 50 targets per ensemble) consistently elicited increased activity in 199 ± 9 neurons (“positive followers”, mean ± SEM), representing 2.5 ± 0.1% of the non-targeted population (**Figure S9B**). A significant decrease was observed in 267 ± 14 neurons (“negative followers”), representing 3.4 ± 0.2%. Interestingly, the numbers of both positive and negative followers were similar when stimulating either random or functionally-defined ensembles (**Figures S9C and S9D**). We then investigated the frequency of positive followers recruited by more than one ensemble, and found that it was significantly higher than expected by chance (**Figure 4C**); a substantial fraction of positive followers (18 ± 3.8%) exhibited recruitment by at least four different ensembles (>20-fold greater than the expectation by chance of 0.87 ± 0.3%), and in fact a robust fraction (6.3 ± 1.8%) was even recruited by eight or more different ensembles (expectation by chance: 0%).

We then analyzed how stimulation-evoked responses changed with distance to the nearest target (**Figure 4D**). When considering the activity averaged across all non-targeted neurons, recruited activity was increased only within 30 µm of the stimulation target and was suppressed at larger distances (consistent with previous studies^20,21^). In contrast, when considering the activity of GER neurons (hereafter defined as positive followers recruited by ≥4 non-overlapping ensembles), activity was remarkably increased even when the closest target was >150 µm away. Interestingly, we found that the recruitment of GER neurons was strongly dependent on stimulation frequency (**Figure S9E**). Together, these results reveal a subpopulation of neurons that exhibit surprising and distinctive recruitment behavior, detectable only with reliable large-scale access to many hundreds of individually-defined cells, that consistently and strongly respond to a diverse range of non-overlapping stimulation patterns, in stark contrast to the other responsive cells.

### GER neurons exhibit distinctive functional properties

To assess whether the distinctive recruitment pattern of GER neurons was linked to specific functional characteristics outside the stimulation context, we analyzed the visually evoked responses of GER neurons from imaging sessions that were acquired the day prior to holographic stimulation experiments (**Figure 5A**, n = 8 mice). We found that GER neurons were visually responsive but untuned for stimulus orientation during drifting grating presentations (**Figures 5B and 5C**). The median fraction of visually responsive neurons that were orientation-selective was significantly lower in GER neurons (0.024) compared to all neurons (0.45; Wilcoxon signed-rank test, p = 0.0078). The response profile to natural scenes was also distinct for GER neurons compared to the remaining population. Unlike the sparse responses we encountered in oScarlet+ neurons in our dataset (e.g. see activity evoked by natural scenes in a representative ensemble of targeted cells, **Figure 3H**), which are typical for excitatory populations in V1 L2/3^40^, GER neurons responded ubiquitously to natural scenes (**Figure S10A**), leading to a significantly reduced sparseness of natural scene responses in these neurons compared to the rest of the population (**Figure 5D**, Wilcoxon signed-rank test, p = 0.0078; response sparseness defined following reference^41^, **STAR Methods**). Thus, the high generality of the GER subpopulation, discovered from responses to large-scale stimulation ensembles, extends also to drifting gratings and natural scenes.

**Figure 5.**
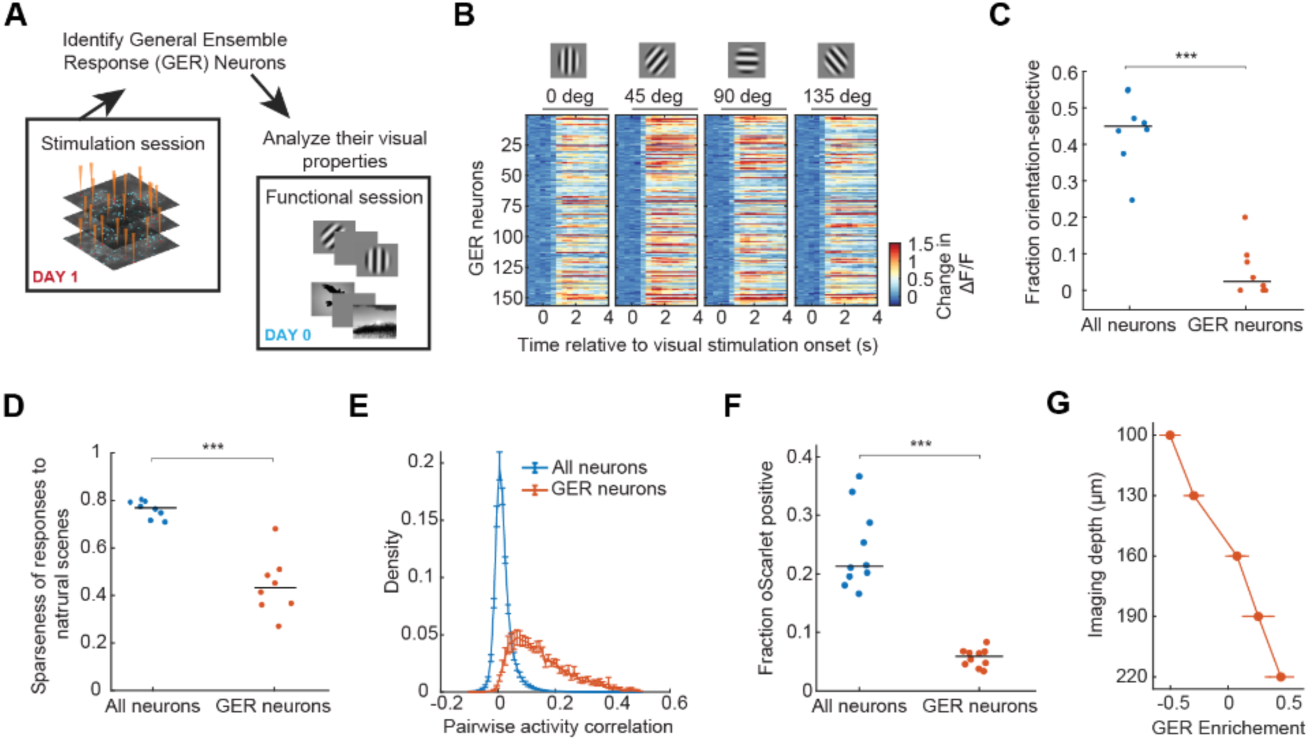
General ensemble-response neurons exhibit distinctive functional properties. **A**, Schematic of the analysis flow. The properties of GER neurons (identified from stimulation sessions) were analyzed based on data from previously acquired imaging sessions with visual stimulation (n = 8 mice). **B**, Trial-averaged change in activity in response to the presentation of gratings with four different orientations for the GER neurons identified in a representative mouse (n = 156 GER neurons). **C**, Fraction of orientation-selective neurons (OSI >0.2) in all neurons and GER neurons. Data points represent values per mouse (n = 8 mice), black bars represent the median across mice. The fraction of orientation-selective neurons was significantly lower in GER neurons (Wilcoxon signed-rank test, p = 0.0078). Neurons non-responsive to grating stimuli were excluded. **D**, Sparseness of the responses to natural scenes (computed on trial-averaged responses and defined following reference^41^, **STAR Methods**) for all neurons and GER neurons. Responses sparseness was significantly reduced in GER neurons (n = 8 mice, Wilcoxon signed-rank test, p = 0.0078). Neurons non-responsive to natural scene stimuli were excluded. **E**, Average histograms of the pairwise activity correlation values of ΔF/F traces during natural scene stimulation for all neurons (left) and GER neurons (right), presented as mean ± SEM for n = 8 mice. The correlation was significantly higher among GER neurons compared to the remaining population (Mann-Whitney U test, p = 1.6 x 10^-4^). **F**, Fraction of oScarlet+ neurons for all neurons (left) and for GER neurons (right) in *Ai230* mice crossed to excitatory Cre driver mice (n = 10 mice). The fraction was significantly higher in all neurons compared to GER neurons (Wilcoxon signed-rank test, p = 0.002). **G**, Relative enrichment of GER neurons across imaging depths (mean ± SEM, n = 10 mice). GER neurons were significantly enriched in the deepest imaging plane (∼220 µm from surface) compared to the most superficial plane (∼150 µm from surface; Wilcoxon signed-rank test, p = 2.0 x 10^-3^).

In addition to their distinct response properties to visual stimuli, pairwise activity correlation during visual stimulation was significantly higher among GER neurons compared to the rest of the population (**Figures 5E and S10B**). This elevated correlation was not simply a consequence of increased general responsiveness to visual stimuli, as shuffling trials substantially diminished the pairwise correlation (**Figure S10C**). Notably, this high degree of correlated activity among GER neurons persisted even in the absence of visual stimulation (**Figure S10D**), raising the possibility that these neurons are intrinsically coupled and form functional subnetworks.

The low degree of orientation tuning^42,43^ in the GER population, as well as their reduced sparseness and high synchrony of natural scene responses^40^, suggested to us that GER neurons could be inhibitory neurons. Although this initial preparation (the Cre-dependent ChRmine-oScarlet reporter *Ai230* crossed to an excitatory Cre driver) did not allow for high-confidence identification of inhibitory neurons due to high background signal in the red channel, which may obscure oScarlet expression, we could confidently identify neurons as excitatory when exhibiting definite oScarlet labeling (**Figure S10E**). We found that GER neurons were much less likely to be oScarlet+ compared to the overall population (**Figure 5F**), indicating that at least a high fraction would be inhibitory neurons. In addition, we found GER neurons to be disproportionately located in deeper imaging planes (**Figure 5G**), which here correspond to deep L2/3. Thus, GER neurons—identified based on their general large-scale ensemble response to holographic stimulation—exhibit specific functional properties during visual stimulation and are likely an inhibitory neuron population present in deep L2/3.

### General ensemble recruitment of inhibitory neurons is unexpected

To date, holographic stimulation of cortical ensembles has been paired with readout of excitatory, but not inhibitory, neurons. To test whether such generalized recruitment of inhibitory neurons upon stimulation of excitatory neurons would be expected given the known connectivity patterns between excitatory and inhibitory neurons in L2/3 mouse visual cortex, we developed a simplified probabilistic network model (**Figures 6A and 6B; STAR Methods**). The model was designed to estimate the probability that a non-targeted neuron would be recruited by any single stimulated ensemble, and thereby calculate the number of ensembles it would be expected to follow, to test the likelihood of existence of GER neurons.

**Figure 6.**
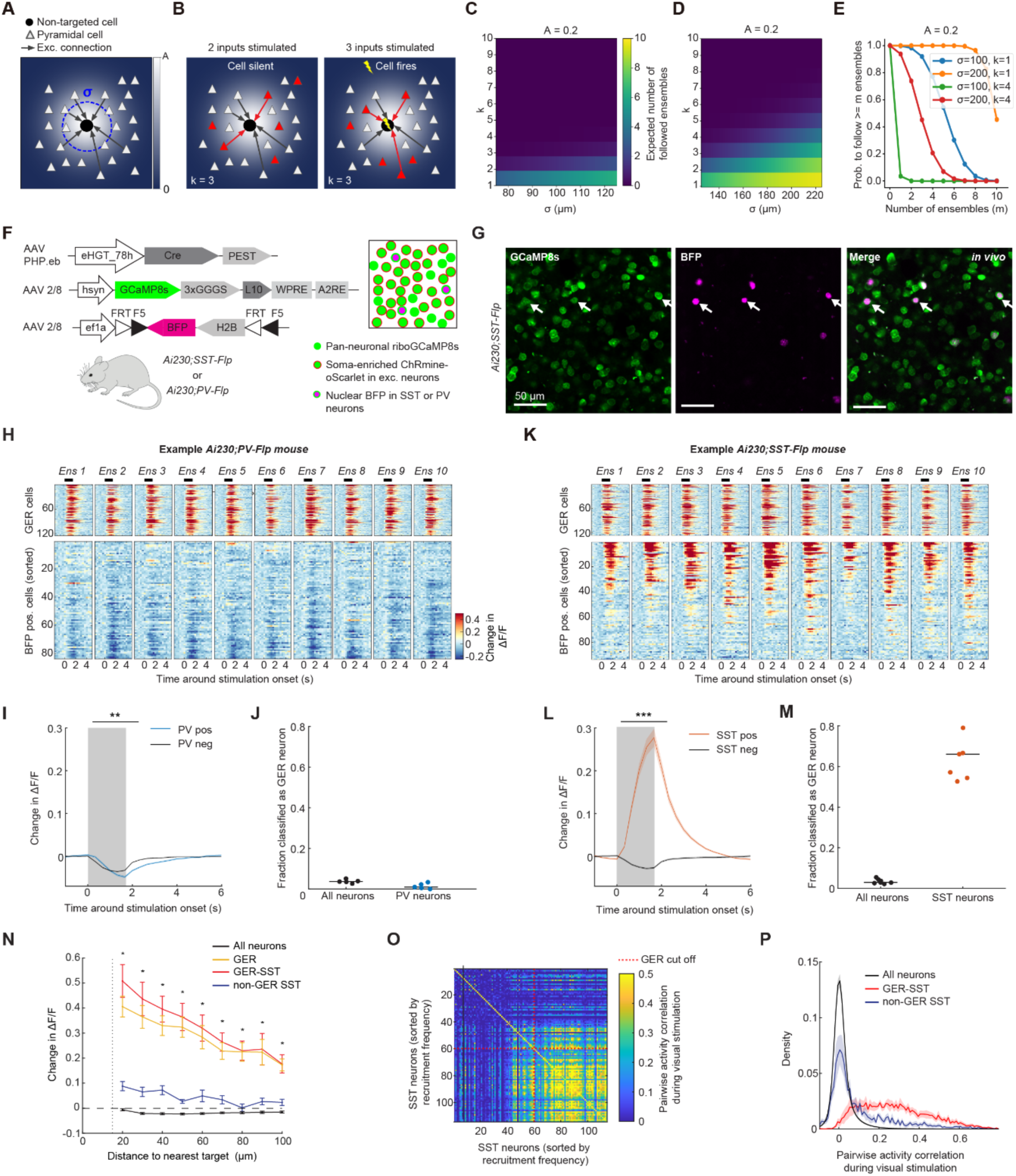
General ensemble-response neurons represent a specific subclass of somatostatin-positive neurons. **A**, Schematic of a simplified probabilistic network model used for estimating non-targeted neuron recruitment probability. The three-parameter model of a non-targeted neuron uses a Gaussian connectivity profile (connection probability indicated by the background color), with peak connection probability A at the origin, and spatial spread σ. **B**, The third parameter of the model is the activation threshold (k), defined as the minimum number of stimulated connected pyramidal cells required to induce firing in the non-targeted neuron. Example: k = 3, firing only occurs with ≥3 stimulated connected cells. **C**, Heatmap showing the expected number of ensembles (out of 10, each with 50 targets) that a non-targeted neuron will respond to, as a function of the activation threshold (k) and spread of connectivity (σ), for a peak connectivity (A) of 0.2. The peak connectivity value and range of σ (75-125 µm) were based on experimental data from pyramidal-to-inhibitory neuron recordings^44^. **D**, Heatmap (as in **C**) for larger σ (125-225 µm). The results indicate that while a larger σ increases the expected number of followed ensembles, a low k value remains necessary to follow a high number of ensembles. **E**, Probability of following ≥m ensembles (out of 10, A = 0.2) for various k and σ. Consistent with panels **C** and **D**, achieving a high probability of following many ensembles (large m) requires large σ and small k. **F**, Schematic of strategy used to add cell-type-specific labeling for inhibitory neurons to the follower-focused approach. *Ai230* mice were crossed with either *PV-Flp* or *SST-Flp* mice, which express Flp recombinase in parvalbumin-(PV) or somatostatin-positive (SST) neurons, respectively. Double-positive mice were systemically injected with AAV-PHP.eb eHGT_78h-Cre-PEST to induce ChRmine-oScarlet expression (eHGT_78h is a pan-excitatory enhancer^32^), followed by intracranial co-injection of two AAVs for hsyn-driven riboGCaMP8s and Flp-dependent nuclear BFP expression. **G**, Representative *in vivo* 2P images in L2/3 visual cortex of an *Ai230*;*SST-Flp* mouse injected with the AAVs shown in **F**. Left: GCaMP8s; Middle: Nuclear BFP; Right: Merged image. Arrowheads indicate examples of co-labeled SST neurons. **H**, Trial-averaged change in activity of GER neurons (top) and non-targeted PV neurons (bottom) in response to stimulation of 10 random ensembles of excitatory neurons (columns, 60 targets per ensemble) in an example *Ai230*;*PV-Flp* mouse. PV neurons were sorted by their mean response across all ensembles (consistent sorting across columns). **I**, Mean stimulation-evoked responses for non-targeted PV neurons (blue) and the remaining non-targeted population (black) during stimulation of random ensembles (shading indicates SEM, n = 50 ensembles from 5 *Ai230*;*PV-Flp* mice). To account for the variability in spatial distribution of targeted neurons across mice, for each mouse we considered only BFP+ neurons within the area encompassing the neurons classified as GER neurons (**STAR Methods**). Responses of non-targeted PV neurons were significantly smaller compared to the remaining non-targeted population (Wilcoxon signed-rank test, p = 9.8 x 10^-3^). Black bar: time window used for quantification. **J**, Fraction of all neurons (black, right) and of PV neurons (blue, left) that were classified as GER neurons. Dots: individual mice. Black bars: median. **K**, Same as **H**, but for GER neurons (top) and non-targeted SST neurons (bottom) in an example *Ai230*;*SST-Flp* mouse. **L**, Same as **I** but for non-targeted SST neurons (red) and the remaining non-targeted population (black). Responses of non-targeted SST neurons were significantly higher compared to the remaining non-targeted population (n = 70 ensembles from 7 *Ai230*;*SST-Flp* mice; Wilcoxon signed-rank test, p = 3.5 x 10^-13^). **M**, Fraction of all neurons (black, right) and of SST neurons (red, left) that were classified as GER neurons. **N**, Stimulation-evoked activity as a function of distance to nearest stimulation target for all non-targeted neurons (black), all GER neurons (yellow), SST neurons classified as GER neurons (red, GER-SST), and SST neurons not classified as GER neurons (blue, non-GER SST). Data is presented as mean ± SEM (n = 7 *Ai230*;*SST-Flp* mice). GER neurons were defined on a left-out set of ensembles (**STAR Methods**). Wilcoxon signed-rank tests assessed GER SST vs non-GER SST (*p ≤0.05). **O**, Heatmap showing the pairwise correlation of SST neuron activity during visual stimulation with drifting gratings for an example mouse. Neurons are sorted by recruitment frequency (highest recruitment at the bottom and right). Red lines indicate the threshold used to classify neurons as GER neurons (recruited by ≥4 ensembles). **P**, Average histograms of the pairwise activity correlation values during visual stimulation with drifting gratings for all neurons (black), GER-SST neurons (red), and non-GER SST neurons (blue) (mean ± SEM across 7 mice).

We developed a three-parameter model of a non-targeted neuron that incorporates a Gaussian spatial profile for the connection probability between any principal excitatory cell and the non-targeted cell. The Gaussian parameters were the peak connection probability A and the spread of connectivity σ (**Figures 6A and S11A**), which we set based on experimental data for pyramidal-to-inhibitory neuron connectivity (A = 0.2, σ = 75-125 µm, see reference^44^). The third parameter, k, represented the activation threshold of the non-targeted neuron, defined as the minimum number of stimulated connected pyramidal cells required to trigger firing in the non-targeted neuron (**Figure 6B**). Given the variable number of active inputs needed to drive firing in cortical neurons, we explored a range of k values (1-10) in our simulations. Using a standard pyramidal-to-inhibitory connectivity profile in mouse cortex^44^, for A = 0.2 and σ = 100 µm (see reference^44^) and activation threshold k = 5, the expected number of followed ensembles was very low (0.004 out of 10), suggesting that the general ensemble-response property in inhibitory neurons is highly unexpected (**Figure 6C**). This low expectation persisted even when considering much higher peak connectivity (A = 0.6) (**Figures S11B and S11C**).

### GER neurons represent a specific class of somatostatin-positive neurons

To understand what could make an inhibitory neuron susceptible to general ensemble recruitment, we used the model to explore different neuronal properties. We found that a low activation threshold (low k) and a large spatial integration range (large σ) both powerfully increased the likelihood of GER properties, as these parameters significantly increased both the recruitment probability of a non-targeted neuron for a given ensemble and the expected number of ensembles followed (**Figures 6D and 6E**). These estimations allowed us to formulate hypotheses to test regarding GER cell typology.

Previous work has shown that at least a subset of somatostatin-positive (SST) interneurons is more readily activated by L2/3 pyramidal neurons compared to parvalbumin-positive (PV) neurons^45–47^. Furthermore, recording SST neurons during optogenetic stimulation of L2/3 pyramidal neurons with increasing light spot sizes revealed that SST neurons integrate over a spatial scale extending 500 µm^45^. Combined with the model, these findings suggested that at least a subset of SST neurons might exhibit some of the requisite properties for the surprising GER phenotype – in particular high susceptibility of firing to very few excitatory inputs, as well as extended spatial integration.

To test experimentally if GER neurons correspond to a specific inhibitory cell type, we added a BFP-label for either one of the two main classes of cortical inhibitory neurons, PV and SST neurons, to the follower-focused approach (**Figures 6F and 6G**). We performed holographic stimulation of random ensembles of excitatory neurons (32 Hz stimulation for 1.5 s, 10 ensembles per mouse, 60 targets per ensemble) and analyzed evoked activity in the non-targeted neurons (n = 5 *Ai230*;*PV-Flp* mice, n = 7 *Ai230*;*SST-Flp* mice). The fraction of neurons classified as GER neurons (positive followers recruited by ≥4 ensembles) was not significantly different between the two mouse lines (Mann-Whitney U test, p = 0.63). To account for spatial target distribution differences across mice, only BFP+ neurons within the area encompassing all GER neurons were considered.

PV neurons were significantly suppressed by large-scale ensemble stimulation (**Figures 6H and 6I**; n = 50 ensembles in 5 mice, Wilcoxon signed-rank test, p = 1.5 x 10^-9^). The magnitude of this suppression decreased as a function of distance to the nearest target (**Figure S10D**). As a consequence of this recruitment pattern, PV neurons were rarely classified as GER neurons (**Figures 6J and S11E**). In fact, the suppression in PV cells was slightly larger compared to the overall population (**Figure 6I**; Wilcoxon signed-rank test, p = 9.8 x 10^-3^), possibly due to elevated background activity in PV cells^45,48^.

In stark contrast to PV neurons, non-targeted SST neurons exhibited increased activity upon large-scale ensemble stimulation (**Figures 6K and 6L**; n = 70 ensembles in 7 mice, Wilcoxon signed-rank test, p = 3.6 x 10^-13^). This increase was driven by a subset of SST cells that strongly responded to multiple ensembles, reminiscent of GER neurons (**Figure 6K**). While less than two-thirds (median 66.1%) of SST neurons in layer 2/3 of visual cortex were classifiable with the GER property (GER-SST neurons, **Figures 6M and S11F**), this fraction was much greater than the 2.9% corresponding to all extracted GCaMP ROIs (n = 7 mice, Wilcoxon signed-rank test, p = 0.016). Moreover, as with purely functionally-defined GER neurons (**Figure 4D**), GER-SST neurons responded to stimulation even when the closest target was 100 µm away (**Figure 6N**). In contrast, SST neurons that were not classified as GER cells (non-GER SST neurons, 33.9% of all SST neurons) were significantly less activated than GER-SST neurons even when close to a target (**Figure 6N**), indicating that this large-scale circuit property helps resolve specific subclasses of SST neurons.

Consistent with GER-SST neurons forming a robustly-defined and distinct functional class of SST neurons, visually-evoked activity among GER-SST neurons was highly correlated during presentation of drifting gratings, unlike the activity of non-GER SST neurons (**Figures 6O and 6P**; n = 7 mice, Wilcoxon signed-rank test, p = 0.016). Finally, we found that most (53.5%) of GER neurons could be categorized as SST cells (**Figure S11G**), even with a viral SST-labeling approach that will not label all SST cells; notably, the visually-evoked activity of SST-positive and SST-negative GER neurons was highly correlated (**Figures S11H and S11I**), indicative of a functionally coherent cell type and suggesting that the true proportion of SST+ GER neurons is likely to be even higher. Together, these data reveal a specific class of SST neurons with remarkable large-scale circuit-level properties, definable via stable causal access to ensembles spanning hundreds to thousands of individually-defined neurons. This need for large-scale but highly-resolved circuit-activity access in order to identify GER neuron typology may implicate these cells in comparably complex and integrative roles, as discussed below.

## DISCUSSION

In this study, we develop and employ a large-scale, cellular-resolution optogenetic approach to interact with different functional elements of cortical circuits in awake mice. In the process of stimulating functionally-resolved ensembles of excitatory cells and tracking population responses from thousands of surrounding excitatory and inhibitory cells in visual cortex, we were surprised to identify a cell type with a highly unique recruitment profile. While stimulation of ensembles of ∼50 excitatory cells in L2/3 led to a net suppressive effect in the overall network, a unique population of non-targeted cells was robustly recruited by diverse ensembles. The general recruitment of this cell population occurred at a rate more than 20 times higher than expected by chance, and was present even when the closest stimulated neuron was >150 µm away. By incorporating labels for interneuron classes into our large-scale all-optical approach, we identified these general ensemble-response (GER) neurons as predominantly SST-positive interneurons. This work highlights the power of stable causal access to ensembles of hundreds to thousands of individually-defined neurons, combined with comprehensive population-wide readout of response dynamics, in order to illuminate fundamental principles of cortical organization.

### A new toolkit for all-optical preparations over expanded spatial and temporal scales

Despite the growing power and adoption of all-optical methods, a key technical bottleneck has hindered scientific discovery: the robust generation of high-quality all-optical preparations with co-expression of indicators and opsins within the same cells in brain tissue. Previous strategies struggled to achieve the critical combination of widespread, homogeneous, and stable expression of both components, alongside balanced opsin-to-indicator ratios, soma-enrichment of the opsin for precise targeting, high opsin sensitivity for large-ensemble stimulation, and dense neuronal labeling for single-cell read access to large populations over months.

Our work presents tools that fundamentally advance this approach along multiple dimensions. Users can leverage either fully-transgenic or the follower-focused hybrid viral-transgenic technique, depending on application-specific needs. All methods presented herein rely on transgenic expression of soma-enriched ChRmine. We find that CAG-driven soma-enriched ChRmine expression in the TIGRE locus leads to stable, homogeneous, and widespread expression with minimal variability across animals, at levels well suited for efficient 2P stimulation.

We selected the channelrhodopsin ChRmine^13,28^ for large-scale 2P holographic stimulation due to its exceptionally high potency^14^. While faster-closing opsins offer higher temporal precision for spike train reproduction^14^, ChRmine’s large photocurrents and slower closing kinetics enable the co-activation of more neurons at a given laser power. Minimizing power helps to reduce the heating of the tissue from light absorption, which could lead to thermal damage^24,26^. ChRmine’s high potency is especially advantageous for holographic read-write experiments in large brain volumes, as light delivery efficiency decreases with both depth (due to tissue scattering) and distance from the center (due to SLM properties^49^). Furthermore, ChRmine’s potency translates to a high success rate in driving targeted neurons—crucial for activating specific, potentially sparse, neuronal ensembles—reliably across trials, and allows for stimulation periods of ≤2 ms. This short duration minimizes data loss due to imaging detector saturation during stimulation (**Figure S2A**), which is vital for investigating the simultaneous, stimulation-induced effects in non-targeted neuronal populations. Finally, ChRmine’s potency permits minimizing opsin expression, thereby reducing cellular burden.

### Robust fully-transgenic all-optical strategies with high sensitivity

These new fully-transgenic strategies offer a convenient and robust way to generate all-optical preparations with widespread expression of a highly potent opsin and cytosolic indicator, complementing existing fully-transgenic all-optical strategies^25^. In most cases *Ai228* will be preferable to *Ai229* (unless the available red channel in *Ai229* is specifically needed for additional labeling), primarily due to *Ai228*’s superior cross-stimulation performance. In L2/3 of cortex in *Ai228*, high-quality functional signals are obtainable with 20-30 mW imaging laser power, which is far below the detectable cross-stimulation threshold in a 3-plane 1x1 mm^2^ FOV experiment (**Figure S3**). Additionally, the presence of an IRES2 sequence in *Ai228*’s CAG-tTA cassette likely results in lower tTA levels compared to *Ai229*, potentially enhancing long-term cell viability.

While our characterization focused on cortical experiments, *Ai228* is likely suitable for all-optical interrogation of other brain regions, provided effective Cre delivery strategies are identified for those regions. In our experience, finding suitable Cre strategies requires careful selection and often involves a trial-and-error process with parallel testing of multiple approaches (Table S1). Local AAV injections for Cre delivery induced injection site toxicity, even at low titers. Therefore, we generally recommend either employing Cre driver lines (if needed, expression could be down-regulated using doxycycline during development; *Ai228* and *Ai229* both have a Tet-Off design), or, alternatively, systemic administration of blood-brain barrier-crossing AAV variants combined with attenuating Cre function (i.e. by using a PEST sequence that targets the protein for degradation by proteases). We used the pan-excitatory enhancer eHGT_78h^32^ to induce expression in the ChRmine transgenics. Combining the lines with other recently developed cell-type or layer-specific enhancers^50,51^, along with newly developed Cre mutants to maintain cell-type specificity^50^, represents a powerful future approach for obtaining cell-type or layer-specific all-optical preparations.

### Key features of the order-of-magnitude scaling up approach

Our new follower-focused approach enables a more complete view of the downstream effects in the surrounding network than any other all-optical preparation. Combining transgenic ChRmine expression with hSyn-driven, ribosome-tagged^36,52^, expression-enhanced GCaMP8s gives rise to an all-optical preparation with dense, pan-neuronal labeling and dramatically decreased neuropil contamination. The reduced background signal improves ROI detection^52^, enabling activity monitoring of ∼2,000 excitatory and inhibitory neurons in a single 1x1mm^2^ imaging plane, in ChR-GCaMP co-expressing tissue that also features write access. Furthermore, ribosome-tagging is particularly helpful for the detection of downstream activity changes in the non-targeted population because signal contamination due to local GCaMP-labeled neurites of directly photo-stimulated neurons becomes less likely. Thus, the follower-focused approach enables the discovery of new cell types by virtue of their large-scale circuit properties and is particularly well-suited for studying how precisely-defined stimulation patterns alter the dynamics within the dense network of surrounding cortical neurons.

Moreover, this hybrid transgenic-viral approach decouples opsin and indicator expression, allowing for independent control over the targeted cell type and labeling density for each component. We used the excitatory driver *Slc17a7-Cre* to achieve read-write access to a pool of >1,000 neurons within a single acquisition volume. Depending on an application’s need, the hybrid strategy also permits sparsifying ChRmine expression within a cell type or directing it to rarer cell types. Combining sparse opsin expression with dense indicator expression can be a powerful tool for estimating or minimizing the impact of off-target stimulation.

Ultimately, choosing between a fully-transgenic (e.g. *Ai228;Camk2aCreERT2*) and the follower-focused, hybrid transgenic-viral approach depends on application-specific goals. A drawback of the hybrid approach may be its intracranial AAV injection, though in our experience, the ribosome-tagged, expression-enhanced hsyn-driven GCaMP8s construct developed in this study offers remarkable spread and stability compared to cytosolic GCaMP AAVs. Regardless, by significantly expanding both spatial and temporal scales, the new approaches presented herein enable unprecedented cellular-resolution read-write access across brain volumes that not only enhance the robustness and ease of existing experimental designs but also facilitate entirely new types of experimental discovery previously unattainable.

### Unique large-scale recruitment patterns across interneuron classes

Feedback inhibition is a hallmark of local cortical circuits, particularly in L2/3. Reflecting the strong local loops between excitatory and inhibitory neurons, 2P all-optical studies reading and writing activity of L2/3 excitatory neurons reported widespread suppression as the dominating network effect^11,16,20,21^; previous work has also shown a net suppressive effect on excitatory neurons evoked by 1P stimulation of ∼100 excitatory neurons in L2/3^53^. While this effect is thought to be mediated by disynaptic inhibition, inhibitory neurons were not monitored in the 2P cellular-resolution optogenetic studies.

Employing a pan-neuronal readout approach and adding cell-type specific labels to different interneuron classes, we found that holographic 2P stimulation of large-scale excitatory ensembles in L2/3 robustly suppressed PV neurons, however a subclass of SST neurons (SST-GER neurons) became generally activated. These data suggest that, in response to activation of large ensembles of L2/3 principal neurons under our stimulation conditions (∼50 neurons per ensemble; near-simultaneous stimulation of all individually-specified neurons at 32 Hz for 1.5 seconds), a subtype of L2/3 SST neurons, and not PV neurons, are the crucial player in providing L2/3 intralaminar inhibition^45,54^. Due to the distinct synaptic dynamics of PV and SST cells^55^ and the supralinear recruitment in SST neurons^54^, it is expected that the relative contribution of SST and PV neurons in L2/3 intralaminar interactions will depend on the specific spatial and temporal properties of the principal cell activation.

### A functional subnetwork of SST neurons closely tracking L2/3 excitatory drive

Cortical SST neurons are a diverse group of cell types with distinct input and output connectivity^56–58^. L2/3 has been reported to contain at least two functionally distinct SST subtypes^59,60^, which aligns with our observation that SST-GER neurons exhibit unique large-scale circuit properties compared to other SST neurons in L2/3.

Our data suggests that SST-GER neurons track the overall level of excitation in surrounding L2/3 pyramidal neurons, as evidenced by their general recruitment across diverse large-scale L2/3 ensembles with strong dependence on the stimulation frequency. We also found that GER neurons synchronize their activity with each other, but not with non-GER SST neurons, during both visually-evoked and spontaneous activity. Together, this indicates that SST-GER neurons form a functional subnetwork (potentially electrically-coupled^46,59,61^), with its activity able to closely track sustained activation of a pool of surrounding principal neurons. Given the remarkably conserved typology of inhibitory cell types across neocortex^62^, a similar architecture is expected to be present in other cortical areas.

By robustly counteracting sustained excitatory drive through their inhibitory action, SST-GER neurons may act as critical regulators of cortical circuit stability. More specifically, by summing the activity of a pool of L2/3 principal neurons and providing inhibitory feedback to individual principal neurons in L2/3 and L5 (the major outputs of SST neurons), SST-GER neurons could dynamically adjust principal neuron sensitivity inversely with the network’s overall activity level. This operation could equip cortical circuits with the capability to perform normalization, which can yield a variety of functions, including facilitation of robust stimulus discrimination across varying input strengths, and enabling computations invariant to certain stimulus dimensions^63^. Furthermore, GER-SST inhibition might be involved in redundancy reduction, making representation in L2/3 more efficient, as well as adding the capacity for winner-take-all behavior to the circuit.

Given that GER neurons are predominantly L2/3 SST neurons, and L2/3 SST axons mostly target the distal dendritic tufts of pyramidal neurons^55,58^, it is an intriguing question why such a normalizing inhibitory signal would target dendrites and not the soma, as somatic inhibition would be the most effective way to modulate a neuron’s sensitivity, due to its direct influence on the spike initiation zone. Instead, by targeting distal dendrites, GER-SST neurons could achieve compartmentalized gain control, enabling the selective gating of specific inputs like top-down (feedback) information arriving in L1^64,65^, in contrast to a global effect that would also impact local recurrent activity on more proximal dendrites. Thus, while GER-SST neurons are broadly recruited by the activation of L2/3 excitatory neurons, their circuit action might be specialized rather than general. In addition to targeting specific neuronal compartments, it is possible that GER-SST neurons participate in cell type specific circuits by targeting specific cell types or layers^66,67^, as has been described for SST subtypes in L5^56–58^.

### Potential implication in neuropsychiatric disorders

SST cells have been broadly linked across psychiatric disorders. Our data, revealing a unique recruitment pattern of a subclass of SST neurons, prompts questions about the potential role of this cell type in certain pathological brain states. Notably, impaired ability of neural circuits to normalize—as discussed above, a potential consequence of SST-GER cell dysfunction—could give rise to a broad range of cognitive deficits. This includes a reduced dynamic range in sensory, motor, emotional systems, alongside attentional deficits due to diminished winner-take-all competition^63,68–72^.

Remarkably, SST cells are the single cell type most strongly linked to schizophrenia (more so than PV neurons, for example as shown by a study combining single-cell transcriptomics and disease genetics from GWAS screens^73^). Schizophrenic patients exhibit numerous EEG abnormalities linked to cortical dysfunction^74^, and research is increasingly focusing on SST neurons in generating these EEG abnormalities^69,75^. We note that for SST-GER neurons, their unique general recruitment properties, integration over long distances, and high synchrony—together with their presumed strong influence on the dendritic compartment of principal neurons—suggests that these neurons could be key determinants of cortical rhythms and modulators of network oscillations.

Additionally, our data indicate that SST-GER neurons are crucial for setting up L2/3 intralaminar inhibition, especially in response to excitatory drive arriving in L2/3 when not accompanied by external sensory input or simultaneous excitation from coupled circuitry. It is intriguing to speculate that impaired GER-SST function would favor the propagation of internally generated (e.g. top-down) signals arriving in L2/3 by diminishing the powerful local suppressive action of GER-SST cells, a possibility further supported by the presumed dendritic action of GER-SST inhibitory feedback. In this way, abnormal GER-SST function could contribute to increased gain of internally-generated signals versus feed-forward inputs, potentially facilitating the propagation of locally-produced patterns uncoupled to more widely-shared timing dynamics (a scenario that bears relevance to the potential circuit mechanisms of psychiatric disorder-related symptoms such as delusions and hallucinations).

### Limitations of our approach

With the access to thousands of read-write neurons and the ability to generate arbitrary light patterns at high temporal resolution using holographic photostimulation, the parameter space of possible stimulation patterns is expansive. Future studies will be needed to investigate how our findings extend to varying stimulation parameters and resulting activity patterns. For example, varying stimulation frequency or controlling the synchrony of stimulated cells—which is easily achieved with the precise temporal and spatial control enabled by 2P holographic stimulation—is expected to alter the balance of PV and SST cell activation. This is because synapses from pyramidal cells onto SST cells are known to facilitate, causing SST cells to act as integrators that respond most strongly to sustained, high-frequency input. In contrast, synapses onto PV interneurons are depressing, leading them to respond most robustly to initial spikes, but less to sustained activation^55^.

Finally, we note that when utilizing *Ai228* or *Ai229*, it is important to confirm that the cell type of interest is not significantly affected by TRE silencing, a phenomenon observed in populations like GABAergic cortical neurons^29^, as TRE-driven GCaMP6m expression would be compromised in those cells. For such cases, a hybrid transgenic-viral approach independent of TRE-driven expression (e.g. *Ai230* with virally delivered hSyn-driven GCaMP) presents a suitable alternative.

**Figure S1.**
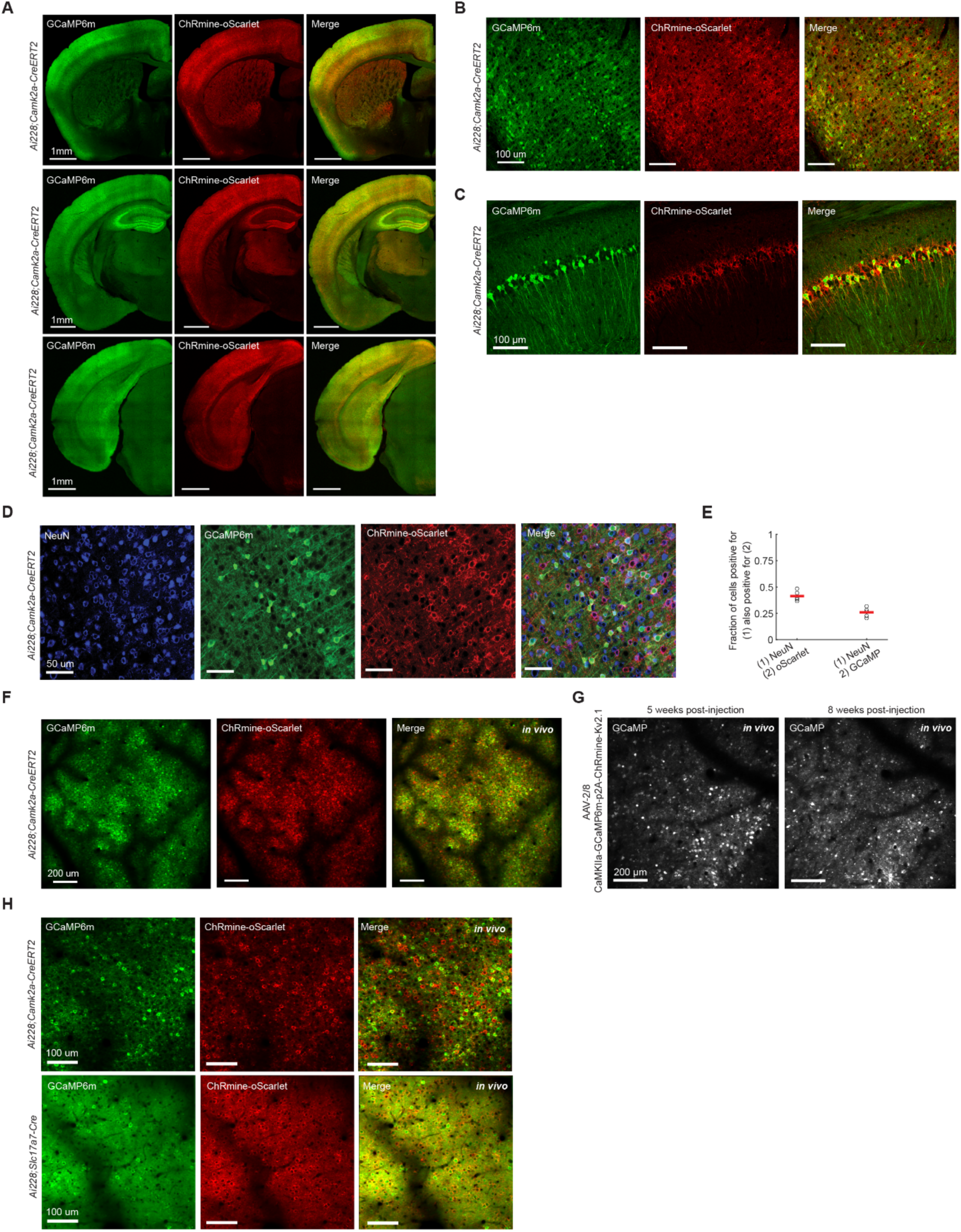
Related to Figure 1. **A**, Confocal images of three coronal sections from a tamoxifen-induced *Ai228*;*Camk2a-CreERT2* mouse, progressing from anterior (top row) to posterior (bottom row). Left column: Native GCaMP6m, middle column: Native ChRmine-oScarlet, right column: merge. **B**, Confocal images of coronal sections of the visual cortex from an *Ai228*;*Camk2a-CreERT2* mouse, showing native expression of GCaMP6m (green) and ChRmine-oScarlet (red). **C**, Confocal images of the CA1 region of the hippocampus from an *Ai228*;*Camk2a-CreERT2* mouse showing native expression of GCaMP6m (green) and ChRmine-oScarlet (red). **D**, Representative confocal images of cortical neurons in a coronal section from an *Ai228*;*Camk2a-CreERT2* mouse stained with the neuronal marker NeuN. Images show NeuN signal (blue), native GCaMP6m (green), and native ChRmine-oScarlet (red). **E**, Quantification of co-labeling of NeuN and oScarlet (left) and NeuN and GCaMP6m (right) in the cortex of *Ai228*;*Camk2a-CreERT2* mice (n = 5 FOVs, 2 mice, example images shown in **D**). Red bars indicate mean values. 41 ± 2.2% (mean ± SEM) of NeuN-positive neurons expressed ChRmine-oScarlet, while 26 ± 2.0% of NeuN-positive neurons expressed GCaMP. **F**, *In vivo* 2P images with a large FOV (1.4 x 1.4 mm^2^) centered over visual cortex in an *Ai228*;*Camk2a-CreERT2* mouse. Green: GCaMP6m. Red: ChRmine-oScarlet. **G**, 2P images of L2/3 visual cortex from a wild-type mouse intracranially injected with AAV-2/8 CaMKIIa-GCaMP6m-p2A-ChRmine-Kv2.1 (3 x 10^11^ vg/ml). The two images show the same FOV at five weeks (left) and eight weeks (right) post-injection. Eight weeks post-injection, previously labeled neurons disappeared, and auto-fluorescent debris accumulated, likely due to cytotoxicity of the construct. **H**, Side-by-side comparison of 2P images from *Ai228*;*Camk2a-CreERT2* (top) and *Ai228*;*Slc17a7-Cre* (bottom) mice. *Slc17a7-Cre*;*Ai228* preparations exhibited higher background GCaMP signal, hindering single-neuron detection. Furthermore, seizures were observed in *Slc17a7-Cre*;*Ai228* mice. Consequently, we discontinued this approach.

**Figure S2.**
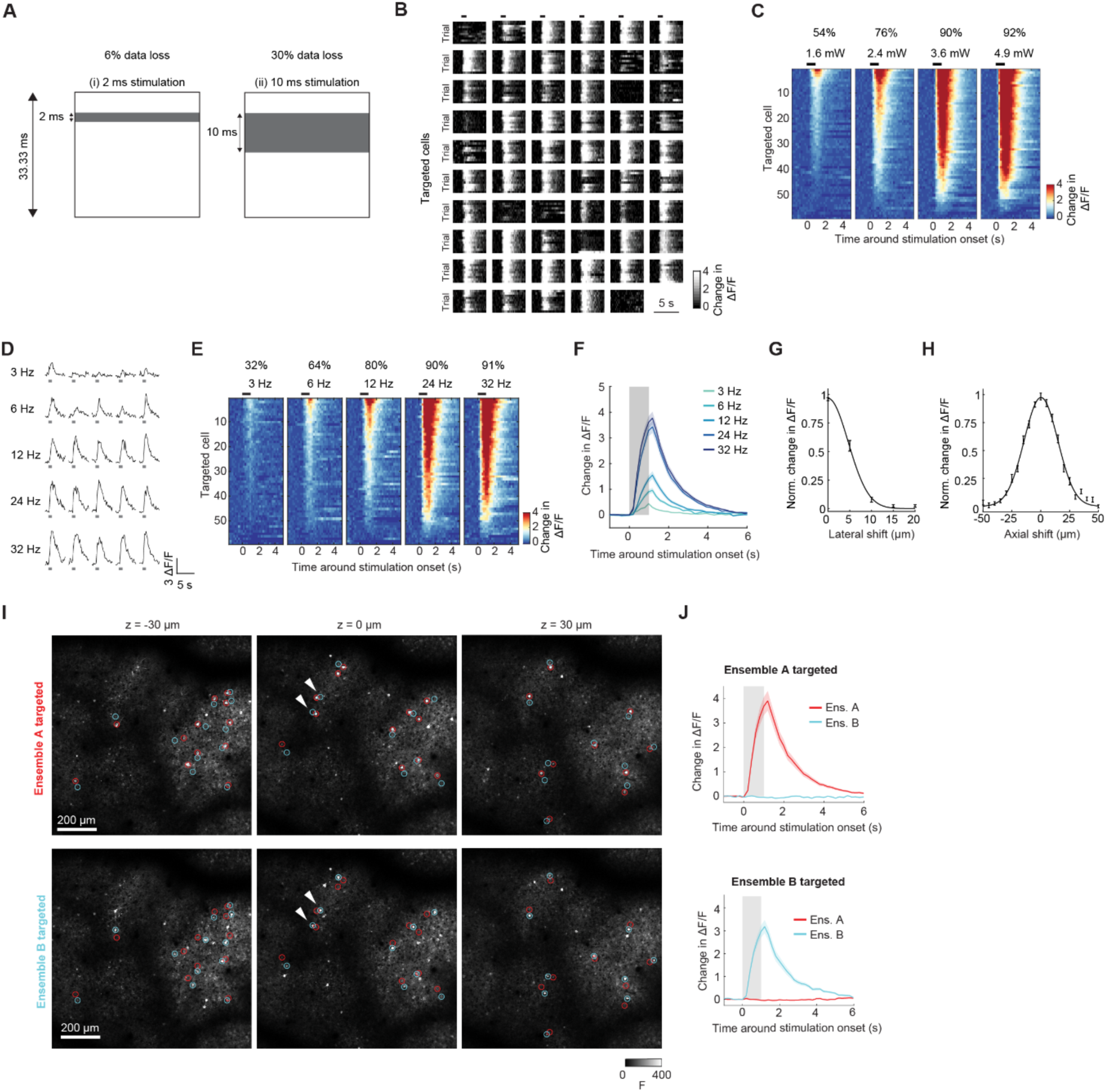
Related to Figure 1. **A**, Schematic illustrating data loss per imaging frame caused by imaging detector saturation during holographic stimulation (“stimulation artifact”) for different stimulation times. An imaging frame (white square) with a 33.33 ms acquisition time (30 Hz imaging rate) is partially obscured by a stimulation artifact (gray). The figure compares the proportion of data loss for a 2 ms (i, 6% data lost) versus a 10 ms (ii, 30% data lost) stimulation period. **B**, Heatmaps of single-trial change in activity in the targeted neurons around stimulation onset (60 targets, 10 trials per power level, 32 Hz, 3.6 mW illumination power/neuron, same ensemble as in Figure 1L). Each plot represents a targeted neuron. Data from targets without an assigned ROI are not shown (**STAR Methods**). Black rectangle on top indicates stimulation duration (1 s). **C**, Trial-averaged change in activity for targeted neurons in response to varying stimulation powers (60 targets, same ensemble as in Figure 1L). Neurons are sorted by response amplitude for each power level. Values on top indicate the percentages of successfully stimulated neurons (**STAR Methods**) for each power level. **D**, Single-trial activity in a representative neuron during stimulation at varying frequencies (3–32 Hz, 3.6 mW illumination power/neuron, 5 trials per frequency). Gray rectangle: 1s stimulation. **E**, Trial-averaged change in activity for targeted neurons in response to varying stimulation frequencies (60 targets, 5 trials per frequency, same ensemble as in Figure 1L). Neurons are sorted by response amplitude for each stimulation frequency. Values on top indicate the percentages of successfully stimulated neurons for each stimulation frequency. **F**, Trial-averaged change in activity averaged across neurons for varying frequencies (same data as in **E**). Shaded area: SEM. **G**, Lateral resolution of the pPSF in L2/3 of an *Ai228*;*Camk2a-CreERT2* mouse, measured by systematically shifting target locations in 5 µm increments. Data shows the average of the normalized ΔF/F (n = 10 trials, n = 37 neurons from 1 mouse; same data in Figure 1Q), fitted with a Gaussian. Error bars represent SEM. **H**, As in **G**, but for axial shifts. Data from 28 neurons from 1 mouse. **I**, Trial-averaged fluorescence images from an example experiment in L2/3 of an *Ai228*;*Camk2a-CreERT2* mouse (3-plane acquisition volume, 30 µm spacing between planes) demonstrating the high spatial precision of the stimulation by targeting two ensembles with closely spaced neuron pairs (Ensemble A: red circles; Ensemble B: cyan circles). Top row: Average fluorescence during Ensemble A stimulation; Bottom row: Average fluorescence during Ensemble B stimulation. White arrowheads highlight two pairs of neighboring neurons, each responding only when directly targeted. **J**, Trial-averaged change in activity averaged across neurons for each ensemble shown in **I** during stimulation of Ensemble A (top) and Ensemble B (bottom). Despite their close proximity, neurons respond only when directly targeted. Shading represents SEM. Pairs of targets with centroid-to-centroid distance <50 µm were included in the analysis (n = 34 pairs, distance between pairs (µm): median 29, range 11-49).

**Figure S3.**
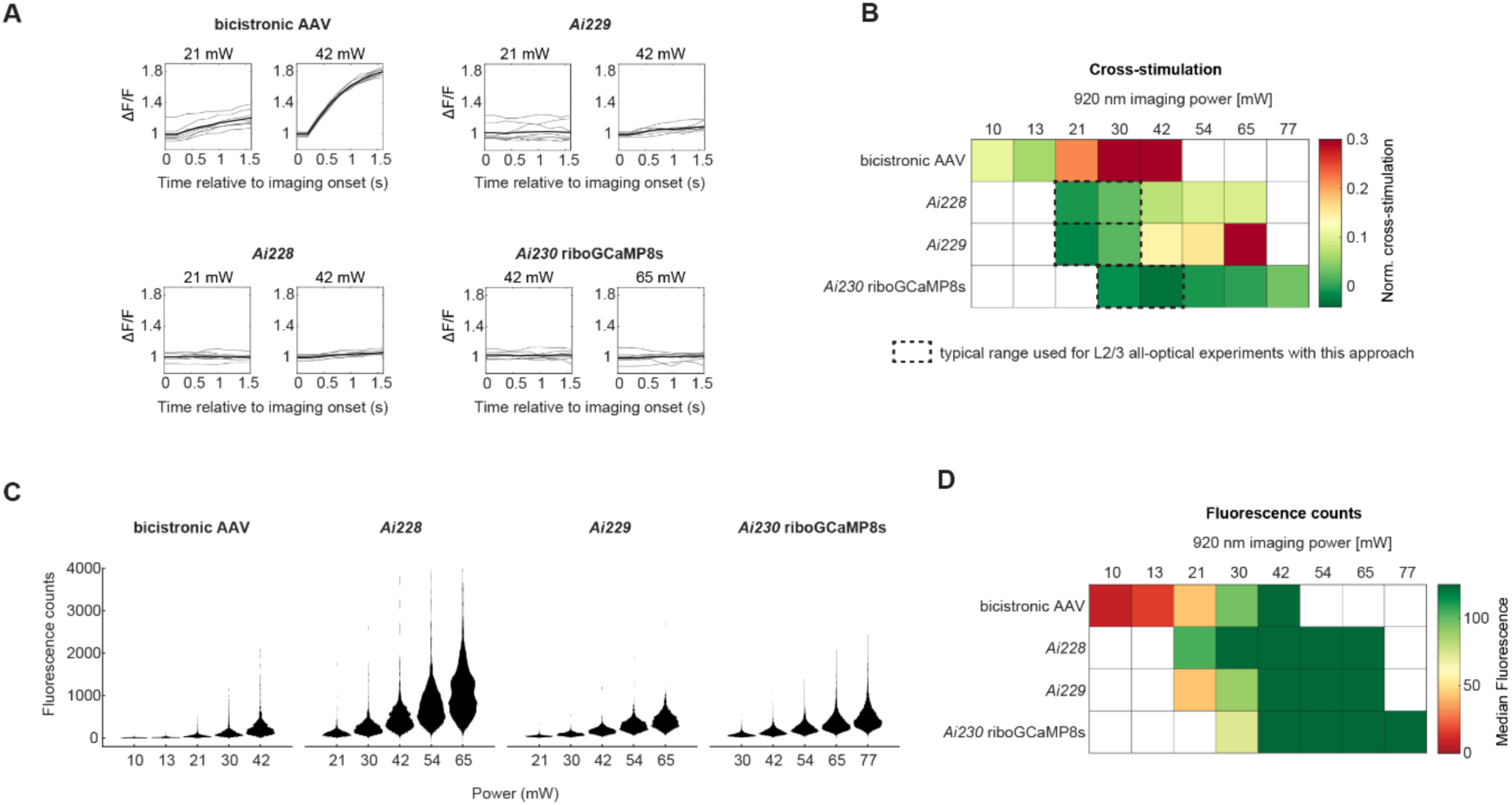
Related to Figure 1. **A**, Representative ΔF/F activity traces during the first 1.5 s of imaging at 920 nm, used to assess unintended ChRmine activation by the imaging laser (“cross-stimulation”). Notably, cross-stimulation will be dependent on the pixel dwell-time of a particular application. Here, data were acquired from L2/3 visual cortex using the following settings: a 3-plane volume with 1 x 1 mm^2^ FOV, 1024 x 1024 pixels per plane, 5 Hz per plane, 30 µm spacing between planes, and an upper plane depth of ∼150 µm. Data represent the average across neurons (gray: single trial data, black: average across trials), normalized by the minimum value of the trial-averaged time series. Data from four different all-optical strategies are shown: “Bicistronic AAV”: wild-type mouse intracranially injected with AAV-2/8 CaMKIIa-GCaMP6m-p2A-ChRmine-Kv2.1 (3 x 10^11^ vg/ml), data acquired 4 weeks post-injection (n = 1,356 neurons from 1 mouse); “*Ai228”*: tamoxifen-induced *Ai228*;*Camk2a-CreERT2* mouse (n = 1,233 neurons from 1 mouse); “*Ai229”*: tamoxifen-induced *Ai229*;*Camk2a-CreERT2* mouse (n = 974 neurons from 1 mouse); “*Ai230* riboGCaMP8s”: *Ai230*;*Slc17a7-Cre* mouse intracranially injected with AAV-2/8 hsyn-GCamP8s-3xGGGS-L10-ARE (n = 3,986 neurons from 1 mouse). **B**, Heatmap quantifying cross-stimulation in the data shown in **A**, including additional imaging power levels. Cross-stimulation was quantified as the difference in trial-averaged ΔF/F between the first and the last frame of the 1.5 s imaging period (**STAR Methods**). White: no data acquired for that approach/power level combination. Dashed black rectangles: typical range of imaging power used in our experiments in L2/3. **C**, Violin plots showing the distributions of ROI fluorescence counts measured at different imaging power levels for the indicated approaches. **D**, Heatmap showing the median of the fluorescence count distributions shown in **C**. White: no data acquired for that approach/power level combination. While the bicistronic viral approach led to significant levels of cross-stimulation at low imaging power levels (21 mW: 0.22 cross-stimulation) where the signal was low (44.2 median fluorescence counts), the transgenic *Ai228*;*Camk2a-CreERT2* approach exhibited minimal cross-stimulation even for high imaging power (<0.09 cross-stimulation for power levels up to 65 mW), while providing high signal already at moderate power (30 mW: 228 median fluorescence counts).

**Figure S4.**
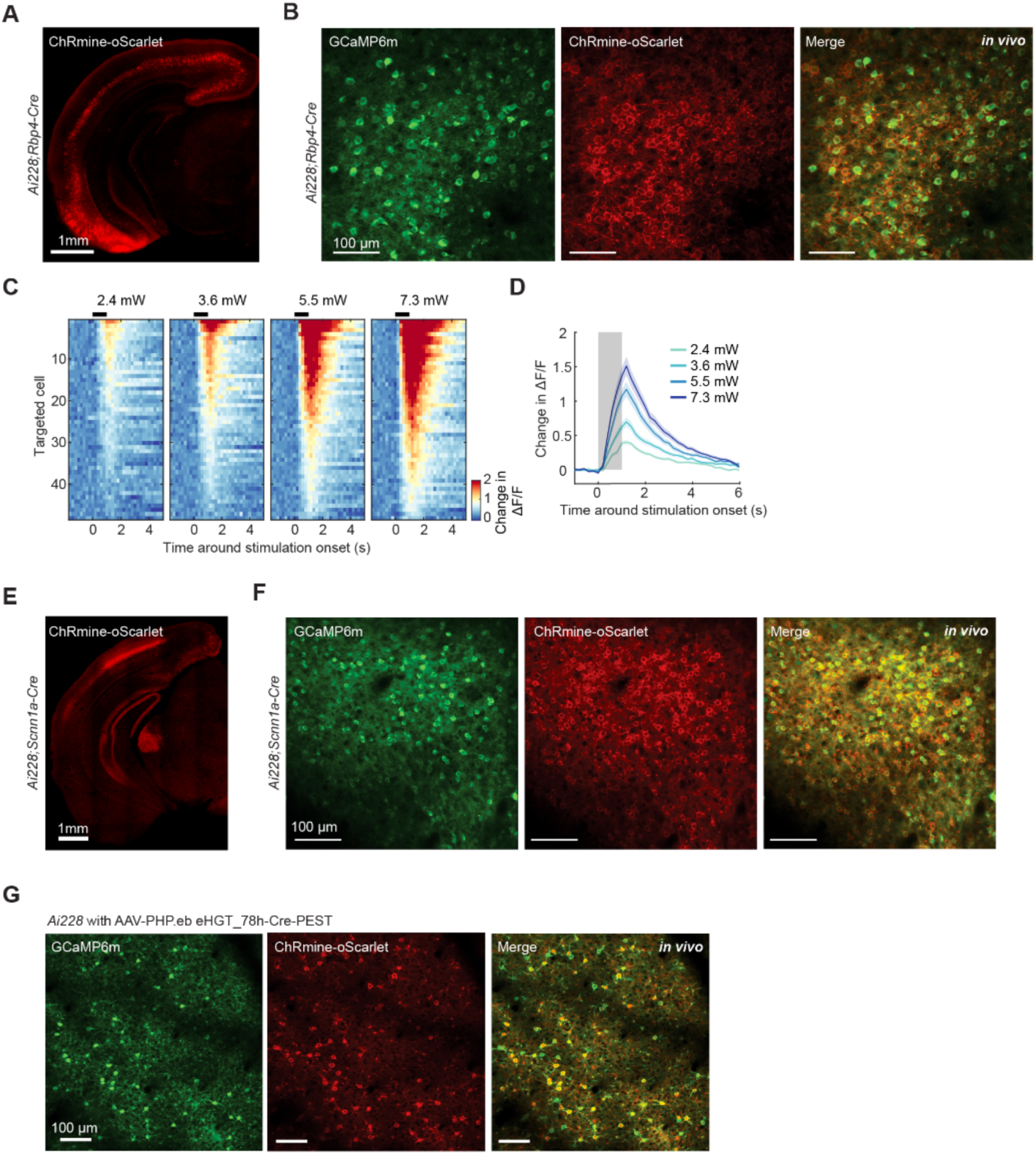
Related to Figure 1. **A**, Confocal image of a coronal brain section from an *Ai228;Rbp4Cre* mouse. In the cortex, soma-enriched ChRmine-oScarlet expression (red) is restricted to L5. **B**, *In vivo* 2P images of an *Ai228*;*Rbp4-Cre* mouse, showing the individual color channels (GCaMP6m: green, ChRmine-oScarlet: red) corresponding to Figure 1R. **C**, Stimulation-evoked, trial-averaged change in activity of targeted L5 neurons in an *Ai228*;*Rbp4-Cre* mouse (48 targets in a 3-plane volume with 30 µm spacing, 1x1 mm^2^ FOV, 1 s stimulation at 24 Hz, 10 trials). Neurons are sorted by response amplitude for each power level. **D**, Trial-averaged ΔF/F activity averaged across neurons (same data as in **C**). Shaded area: SEM. **E**, Confocal image of a coronal brain section from an *Ai228*;*Scnn1a-Cre* mouse. In the cortex, soma-enriched ChRmine-oScarlet expression (red) is restricted to L4. **F**, *In vivo* 2P images of an *Ai228*;*Scnn1a-Cre* mouse, showing the individual color channels (GCaMP6m: green, ChRmine-oScarlet: red) corresponding to Figure 1T. **G**, *In vivo* 2P images of an *Ai228* mouse systemically injected with PHP.eB-coated AAV containing the pan-excitatory enhancer eHGT_78h^32^ and a PEST sequence to attenuate Cre expression (3 x 10^11^ vg/animal).

**Figure S5.**
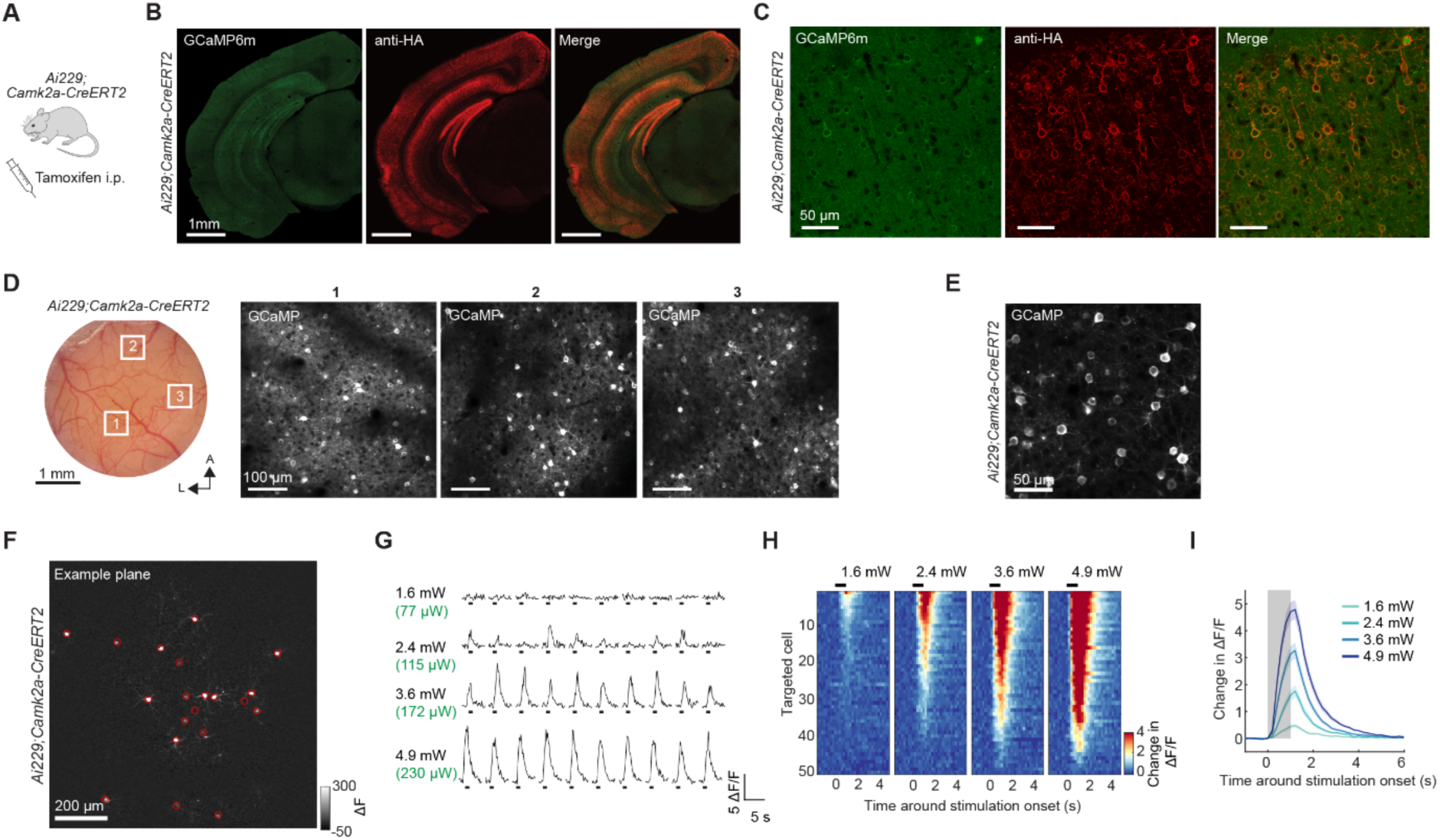
Related to Figure 1. **A**, For all-optical access to excitatory neurons throughout the forebrain, *Ai229* mice were crossed with *Camk2a-CreERT2* mice. **B**, Confocal images of a coronal brain section from a tamoxifen-induced *Ai229*;*Camk2a-CreERT2* mouse. Excitatory neurons throughout the forebrain express GCaMP6m (green) and ChRmine (red, visualized using an HA antibody). **C**, Confocal images of the visual cortex from a tamoxifen-induced *Ai229*;*Camk2a-CreERT2* mouse. **D**, Left: Photograph of a chronic cranial window over the dorsal posterior cortex of a tamoxifen-induced *Ai229*;*Camk2a-CreERT2* mouse. White rectangles indicate the FOVs shown on the right. A: anterior, L: lateral. Right: 2P images showing GCaMP6m expression in L2/3. **E**, Representative 2P image of L2/3 visual cortex in a tamoxifen-induced *Ai229*;*Camk2a-CreERT2* mouse at a higher magnification compared to **D**. **F**, Trial-averaged ΔF image of an example plane during 1s stimulation of 60 simultaneous targets (24 Hz, 3.6 mW illumination power/neuron, 172 µW time-averaged power/neuron, 10 trials, 3-plane acquisition volume). Red circles indicate the targeted neurons (n = 20) in this example plane. **G**, Single-trial activity in a representative neuron during stimulation at varying stimulation powers (10 trials per power level, one power level per row). Values on the left indicate illumination power/neuron (black) and time-averaged power/neuron (green, in brackets). Black rectangles: stimulation period. **H**, Trial-averaged change in activity for targeted neurons in response to varying stimulation powers (60 targets, 10 trials per power level, same ensemble as in **F**). Neurons are sorted by response amplitude for each stimulation power level. **I**, Trial-averaged change in activity averaged across neurons (same data as in **H**). Shaded area: SEM.

**Figure S6.**
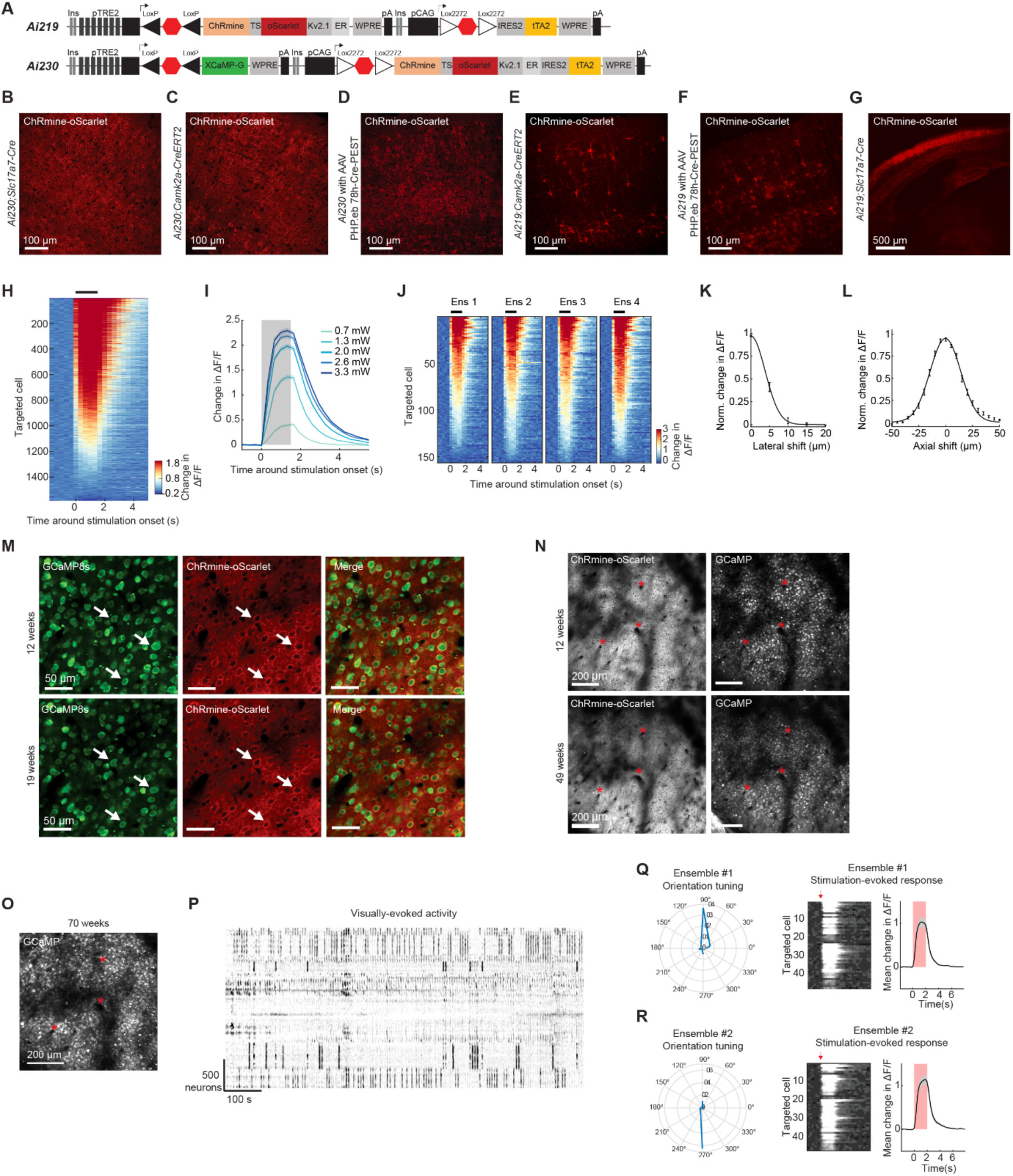
Related to Figure 2. **A**, Schematic diagrams of the new transgenic reporter *Ai219* and the previously presented reporter line *Ai230*^34^. **B**, Representative confocal image of a coronal section of the cortex from an *Ai230*;*Slc17a7-Cre* mouse showing native soma-enriched ChRmine-oScarlet expression (red) in cortical neurons. **C**, Same as **B** but for a tamoxifen-induced *Ai230*;*Camk2a-CreERT2* mouse. **D**, Same as **B** but for an *Ai230* mouse systemically injected with AAV-PHP.eb eHGT_78h-Cre-PEST (3 x 10^11^ vg/animal). **E**, Same as **B** but for a tamoxifen-induced *Ai219*;*Camk2a-CreERT2* mouse. In comparison to *Ai230*;*Camk2a-CreERT2* (shown in **B**), expression levels per cell are higher and expression density is sparser. **F**, Same as **B** but for an *Ai219* mouse systemically injected with AAV-PHP.eb eHGT_78h-Cre-PEST (3 x 10^11^ vg/animal). In comparison to the same Cre delivery approach used in *Ai230* (shown in **D**), expression levels per cell are higher and expression density is sparser. **G**, Representative confocal image of a coronal brain section from an *Ai219*;*Slc17a7-Cre* mouse. This cross resulted in highly abnormal brain morphology with thin cortex and malformed hippocampal formation. **H**, Trial-averaged change in activity of the targeted neurons, pooled across ensembles and ordered by response amplitude, in a representative experiment in an *Ai230*;*Slc17a7-Cre* mouse with ribosome-tagged GCaMP8s (28 ensembles, 60 targets each, same data as in Figure 2F). 1,053 neurons were successfully stimulated. **I**, Trial-averaged change in activity averaged across neurons in response to varying stimulation powers (n = 240 targets, data from 4 ensembles in 1 mouse). Stimulation: 1.5 s, 32 Hz, 10 trials per power level. Values indicate illumination power/neuron. Shaded area: SEM. **J**, Trial-averaged stimulation-evoked change in activity in four ensembles with 160 targeted neurons each. The targets of each ensemble were distributed into two groups, each stimulated with a separate hologram (1.5 s stimulation, 32 Hz, 2.7 mW illumination power/neuron, 5 trials). Stimulation of the entire ensemble occurred within 6 ms. 126 ± 4.6 targeted neurons (mean ± SEM, n = 4 ensembles) were successfully stimulated. **K**, Lateral resolution of the pPSF in L2/3 of an *Ai230*;*Slc17a7-Cre* mouse with ribosome-tagged GCaMP8s, measured by systematically shifting target locations in 5 µm increments. Data shows the average of the normalized ΔF/F (n = 10 trials, n = 28 neurons from 1 mouse; same data in as Figure 2H), fitted with a Gaussian. Error bars represent SEM. **L**, As in **K**, but for axial shifts. Data from 48 neurons from 1 mouse. **M**, *In vivo* 2P images of the same FOV in L2/3 visual cortex of an *Ai230*;*Slc17a7-Cre* mouse intracranially injected with AAV-2/8 hsyn-GCamP8s-3xGGGS-L10-ARE at 12 and 19 weeks post-injection. Arrowheads point to examples of neurons identified across the imaging sessions. **N**, Zoomed-out view of the same preparation shown in **M** at 12 weeks and 49 weeks post-injection. While the preparation was imaged at slightly different angles of head bar and objective, the vessel pattern allows identification of corresponding image locations (red stars). **O**, Same preparation shown in **M** and **N** at 70 weeks post-injection. The vessel pattern allows to identify corresponding image locations (red stars). **P**, Time series of 2,563 neurons in L2/3 visual cortex during visual stimulation with drifting gratings (4 orientations) acquired from FOV shown in **O** (at 70 weeks post-injection). Heatmap shows clustered neuronal activity visualized using ‘Rastermap’^77^. Dark indicates increase in activity. Groups of neurons (“vertical stripes”) are selectively responding to specific grating orientations, showing that >16 months post-injection, orientation tuning is maintained. **Q**, An ensemble of 50 direction-tuned neurons (ensemble #1, tuning shown on the left) was chosen from dataset shown in **P** and targeted for stimulation. Also at 70 weeks post-injection, targeted neurons were stimulated with high efficiency (center: heatmap showing trial-averaged stimulation-evoked response for each neuron, right: mean ΔF/F across neurons). **R**, Same as **Q** but for ensemble #2.

**Figure S7.**
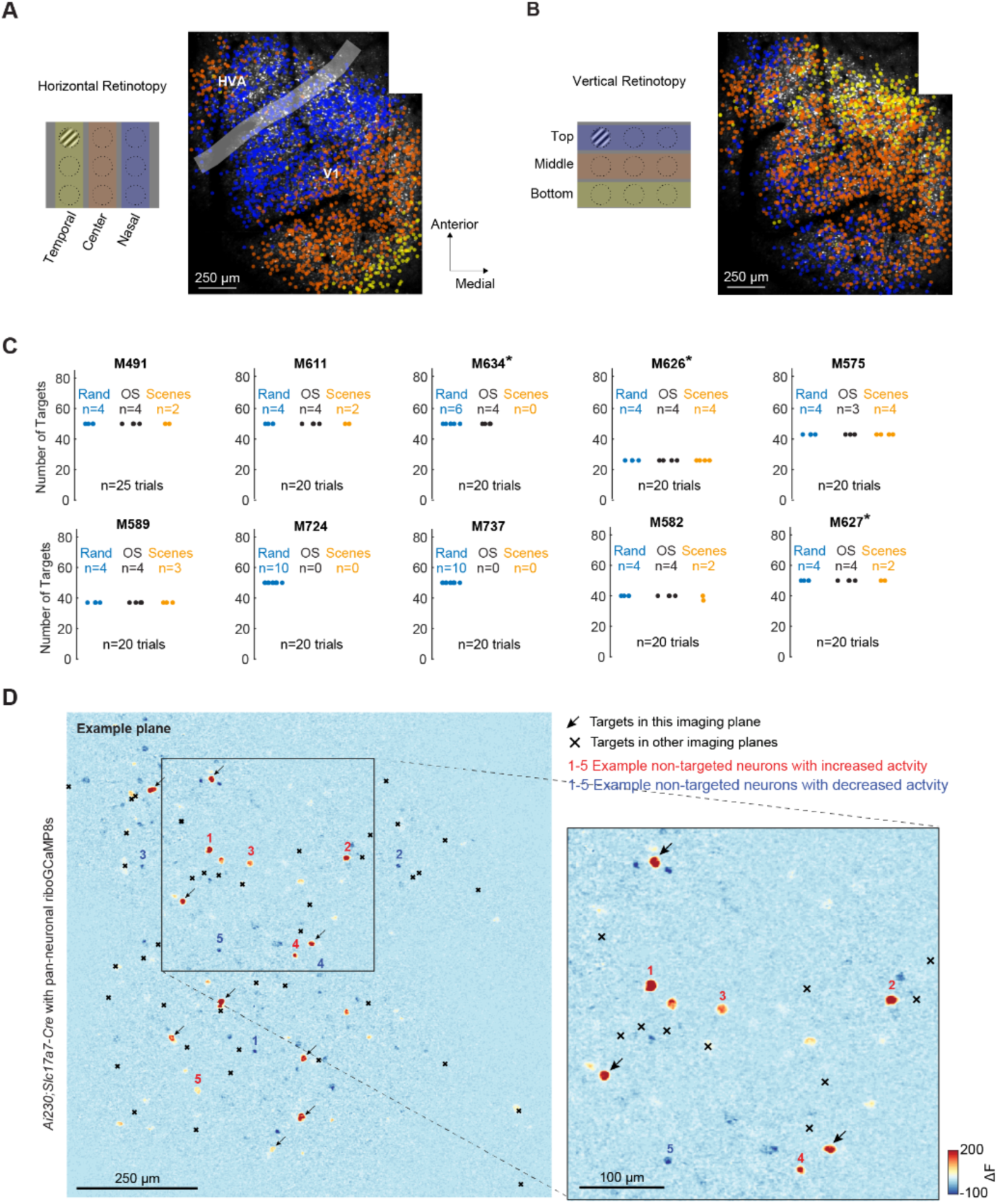
Related to Figure 3. **A**, Retinotopic mapping along the horizontal axis in an example mouse to identify V1 and an adjacent HVA. Left: Schematic of visual stimulation. Small patches of drifting gratings were presented at 9 locations on a screen, indicated by dashed circles. Colors signify the locations of the stimulus (temporal, center, or nasal). Right: 2P images of example planes of two partially overlapping acquisition volumes (each 1380 µm x1380 µm x 4 planes spaced 30 µm apart) that were consecutively imaged during visual stimulation. The positions of neurons, merged across planes, are overlaid. Neurons are color-coded based on their preferred stimulus location along the horizontal axis (temporal, center, or nasal). The border between V1 and HVA was manually delineated by identifying the reversal in the representation of the vertical meridian. For inter-areal experiments, a 1x1mm^2^ FOV was positioned above the V1-HVA region. For V1 experiments, the FOV was positioned above V1. **B**, Same as **A** but for retinotopic mapping along the vertical axis. **C**, Summary of the V1 dataset used in Figures 3, 4 and 5. 7 out of 10 mice were *Ai230*;*Slc17a7-Cre* mice, while 3 mice (marked with an asterisk) were tamoxifen-induced *Ai230*;*Camk2a-CreERT2* mice. The numbers of random (“Rand”), orientation-selective (“OS”), and scene-selective (“Scenes”) ensembles targeted in each mouse are shown. In each mouse, after selection of the orientation-selective and scene-selective ensembles, random ensembles were sampled without replacement from the remaining population of oScarlet+ neurons to create non-overlapping (or disjoint) ensembles (**STAR Methods**). **D**, Trial-averaged ΔF images of an example plane during stimulation of 50 excitatory neurons. Left: Entire plane. Right: Zoom in. Arrows: Targets within this imaging plane (n = 9). Same data as in Figure 3A but also showing the location of the targets of the other four imaging planes (black crosses, n = 41). Locations of the non-targeted neurons that increased activity (i.e. neurons highlighted by red numbers 1-5) were non-overlapping from locations of targeted neurons in any plane, indicating that the co-activation of these neurons is a downstream network effect rather than the result of direct photostimulation.

**Figure S8.**
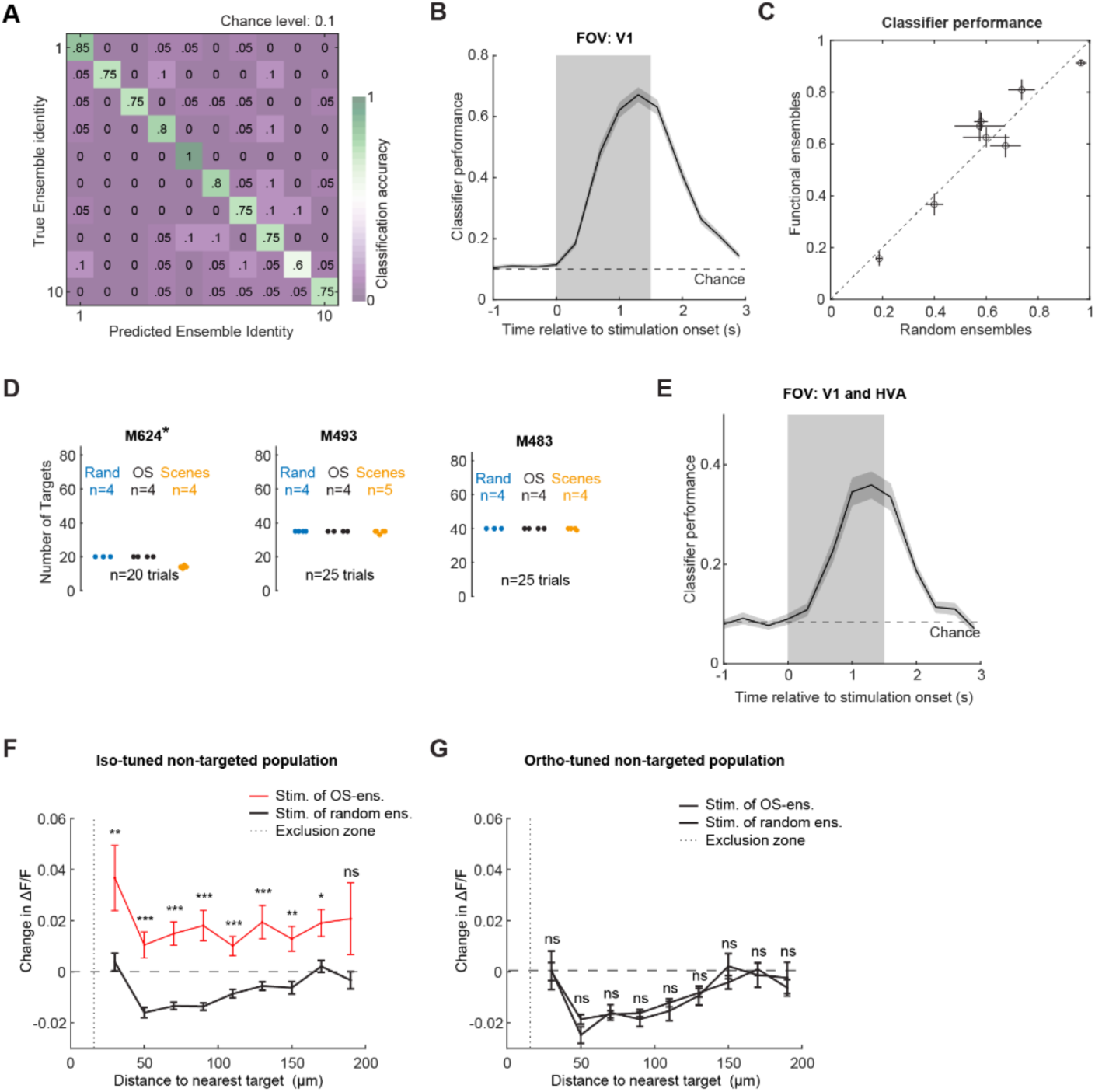
Related to Figure 3. **A**, Confusion matrix from a representative experiment in V1 (10 ensembles, 50 targets per ensemble). The matrix shows the performance of the classifier trained to decode which of the 10 ensembles was stimulated based on the responses of non-targeted neurons. **B**, Sliding window analysis of classifier performance in V1. The trace shows the average classifier performance across all ensembles (n = 104) and mice (N = 10) as a function of time relative to stimulation onset, using 1 time point per window. Shading represents SEM. Dashed line indicates the maximum chance level across mice (0.1, based on 10-12 ensembles per mouse). Gray shading indicates duration of stimulation. **C**, Comparison of classifier performance across ensemble categories. Each point represents the average classifier performance for functional versus random ensembles of a single mouse. Error bars represent the SEM. Dashed line: unity. For quantification, a linear-mixed effects model was fitted to predict the classifier performance based on ensemble identity and other factors (**STAR Methods**). None of the predictors were statistically associated with classifier performance (p>0.05). **D**, Summary of the V1-HVA dataset used in Figure 3. 2 out of 3 mice were *Ai230*;*Slc17a7-Cre* mice, 1 mouse (M624) was a tamoxifen-induced *Ai230*;*Camk2a-CreERT2* mouse. The numbers of random (“Rand”), orientation-selective (“OS”), and scene-selective (“Scenes”) ensembles targeted for each mouse are shown. After selection of the orientation-selective and scene-selective ensembles, random ensembles were sampled without replacement from the remaining population of oScarlet+ neurons to create non-overlapping (or disjoint) ensembles (**STAR Methods**). **E**, Same as **B** but for the V1-HVA dataset (n = 37 ensembles in 3 mice). The dashed line indicates the maximum chance level across mice (0.08, based on 12-13 ensembles per mouse). **F**, Stimulation-evoked ΔF/F responses of iso-tuned non-targeted neurons as a function of distance to the nearest stimulation target when stimulating similarly tuned neurons (red), or a size-matched random ensembles (black). Responses were significantly higher following orientation-selective (OS) ensemble stimulation compared to size-matched random ensemble stimulation for distances up to 170 µm. Data represent mean ± SEM (OS ensembles: 122-1282 neurons per bin; control: 557-5561 neurons per bin; data from 31 OS-ensembles in 8 mice; control: average of 4 random ensembles per mouse; two-sample t-test, Bonferroni-corrected p-values: *** p≤0.001, ** p≤0.01, * p≤0.05, ns p>0.05. **G**, Same as **F** but for the stimulation-evoked responses in ortho-tuned non-targeted neurons, when stimulating orthogonally tuned neurons (blue), or a size-matched random ensembles (black). Responses evoked by stimulation of OS-ensembles were not statistically different from those evoked by random ensembles (OS ensembles: 145-1275 neurons per bin; control: 554-5620 neurons per bin).

**Figure S9.**
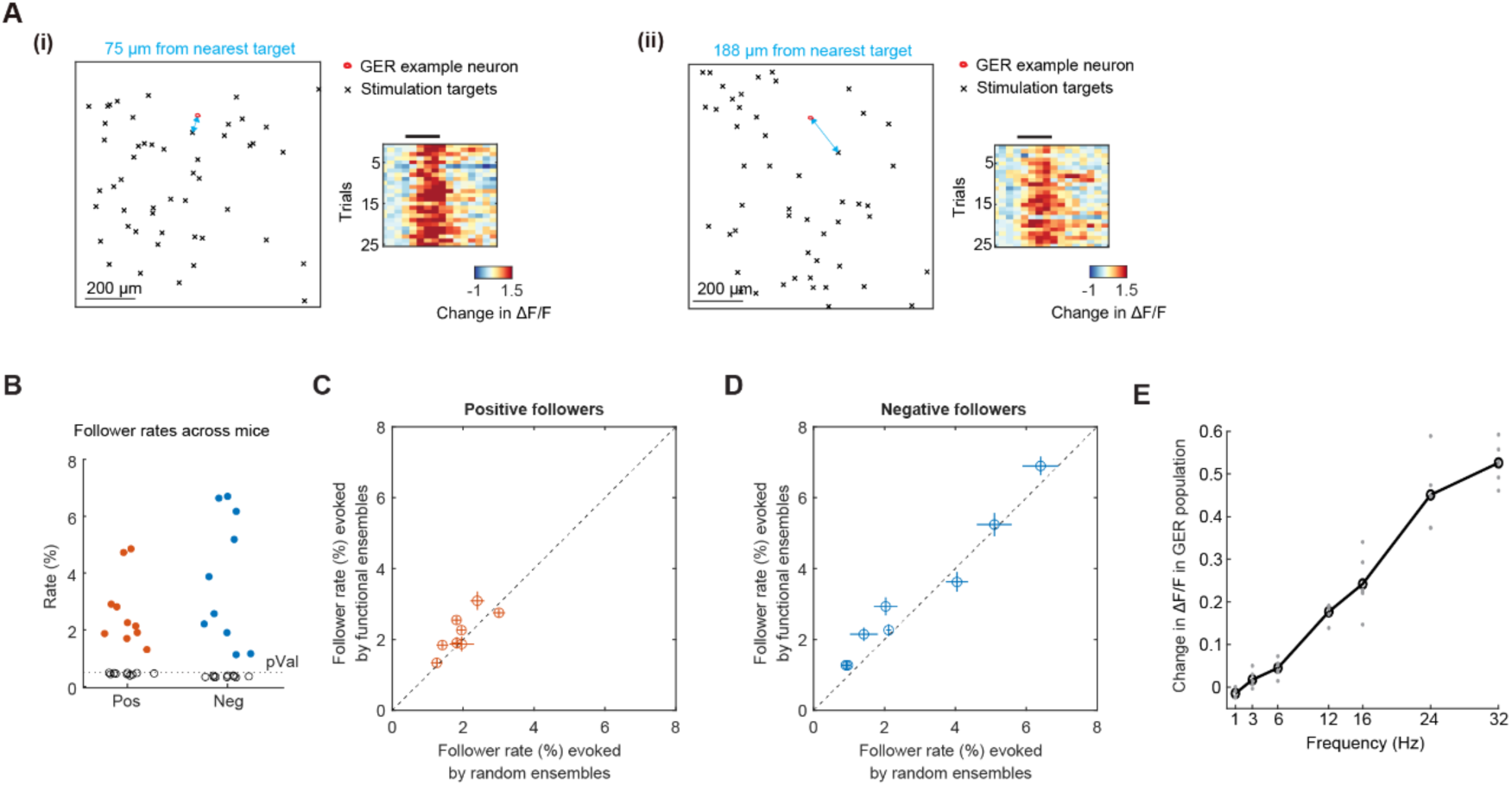
Related to Figure 4. **A**, (i) Left: Location of an example stimulation ensemble (black crosses, locations of 5 imaging planes projected onto a single plane) and an example non-targeted neuron with general ensemble-response (GER) properties (red circle). The distance between the nearest target and the example cell in the 3D imaging volume was 75 µm (blue arrow). Right: Single trial, stimulation-evoked responses of the example neuron during the stimulation of the example ensemble shown on the left. (ii), Same example non-targeted neuron as in (i), but for a different stimulation ensemble (nearest target 188 µm). Black bars indicate the 1.5 s stimulation period. **B**, Percentage of non-targeted neurons (outside of exclusion zone, **STAR Methods**) showing significantly increased (red, positive followers) or decreased (blue, negative followers) activity following stimulation (Student’s t-test p <0.005). Values represent the average across ensembles within each mouse (n = 10-12 ensembles per mouse, N = 10 mice; see also **Figure S7C**). Black circles indicate values obtained from randomly sampled stimulation onsets. The dotted line represents the expected false positive rate (0.5%) based on the t-test significance level. **C**, Comparison of follower rates evoked by the stimulation of functional versus random ensembles. Each point represents the average rate of positive followers across functional and random ensembles for each mouse, lines indicate SEM. Dashed line: unity. For quantification, a linear-mixed effects model was fitted to predict the number of followers (**STAR Methods**). None of the predictors were statistically associated with classifier performance (p >0.05). **D**, Same as **C** but for the fraction of negative followers. **E**, Stimulation-evoked responses of GER neurons as a function of stimulation frequency. GER neurons were defined as non-targeted neurons recruited by ≥4 out of 10 ensembles from a previous stimulation session with 50 targets per ensemble. Gray points show responses averaged across GER neurons to individual ensembles for each frequency. Black curve represents the mean across all ensembles. Data acquired during stimulation of 6 ensembles in 2 mice, 60 targets/ensemble, 1.5 s stimulation at 1-32 Hz.

**Figure S10.**
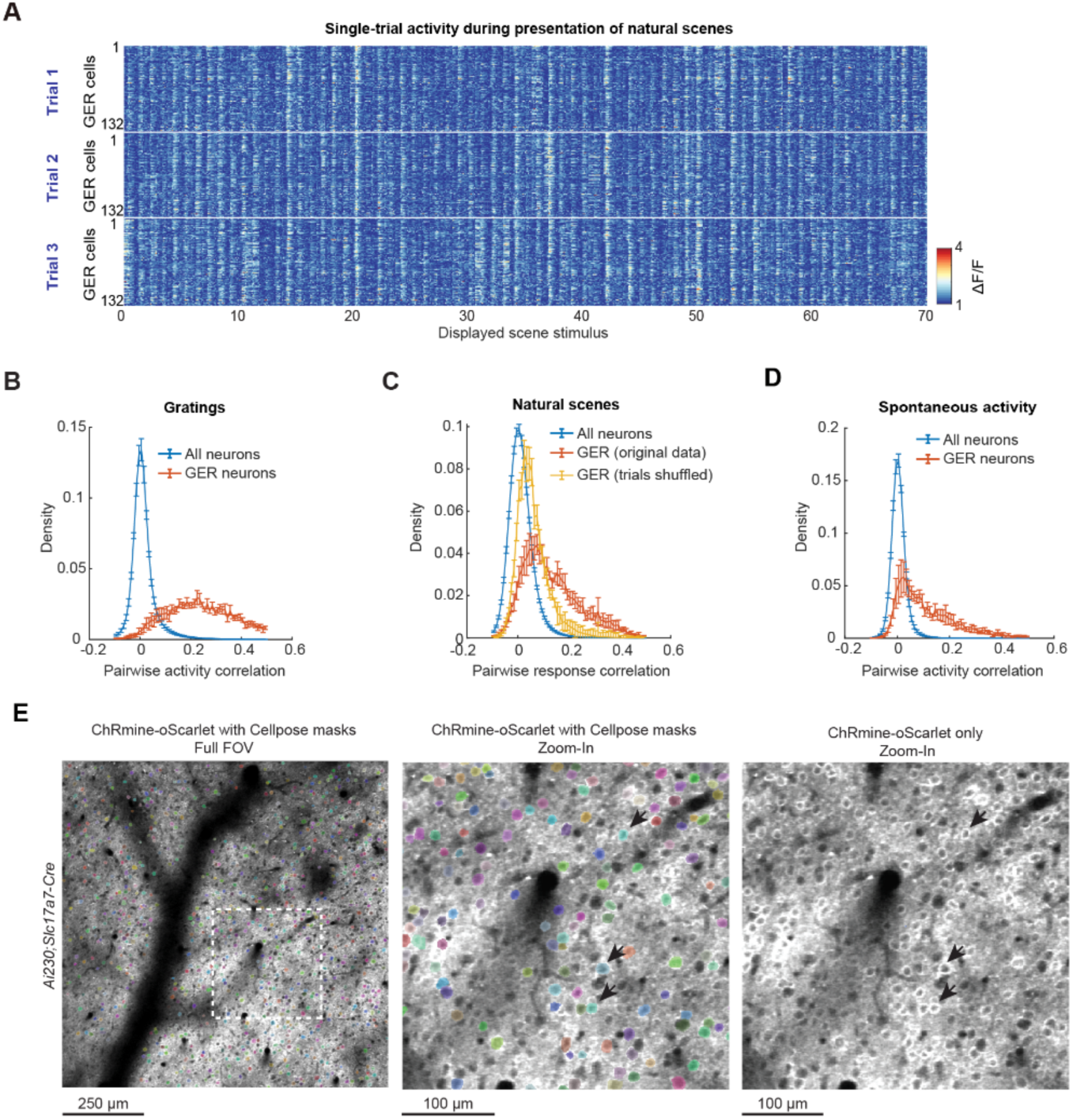
Related to Figure 5. **A**, Single-trial responses of GER neurons to the presentation of 70 natural scene images from a representative mouse. Each of the three blocks shows the responses of 132 GER neurons for one trial, one neuron per row. The sequence of images was repeated for each trial. **B**, Average density of pairwise correlations of activity during presentation of gratings for all neurons (blue) and GER neurons (red, whiskers indicate SEM, n = 8 mice). The correlation was significantly higher among GER neurons compared to the whole population (Mann-Whitney U test, p = 1.6 x 10^-4^). **C**, Average density of pairwise correlation of responses to natural scenes for all neurons (blue) and GER neurons (red: original data; yellow: shuffled trials). Whiskers indicate SEM (n = 8 mice). Shuffling trials significantly reduced the pairwise correlation within the GER neuron population (Mann-Whitney U test, p = 6.2 x 10^-4^), indicating that the elevated correlation among GER neurons (Figures 5E **and S10B**) was not simply a consequence of increased general responsiveness to visual stimuli. **D**, Same as **B** but for activity in the absence of visual stimuli (“spontaneous activity”, last 3 s of the inter-stimulus time interval during holographic stimulation sessions). The correlation was significantly higher among GER neurons compared to the remaining population (Mann-Whitney U test, p = 1.6 x 10^-4^). **E**, 2P images showing ChRmine-oScarlet (gray) and the results of automatic cell segmentation using Cellpose^78^ in a representative plane. Left: full FOV with Cellpose masks superimposed; Center: Zoomed-in region from the left image; Right: Same zoomed-in view, but without Cellpose masks. Black arrows indicate three example oScarlet+ cells. Our segmentation strategy was optimized to prioritize minimizing false positives, ensuring that all identified cells expressed oScarlet. This optimization, combined with considerable background labeling, meant that some oScarlet+ cells were likely not captured by our segmentation method.

**Figure S11.**
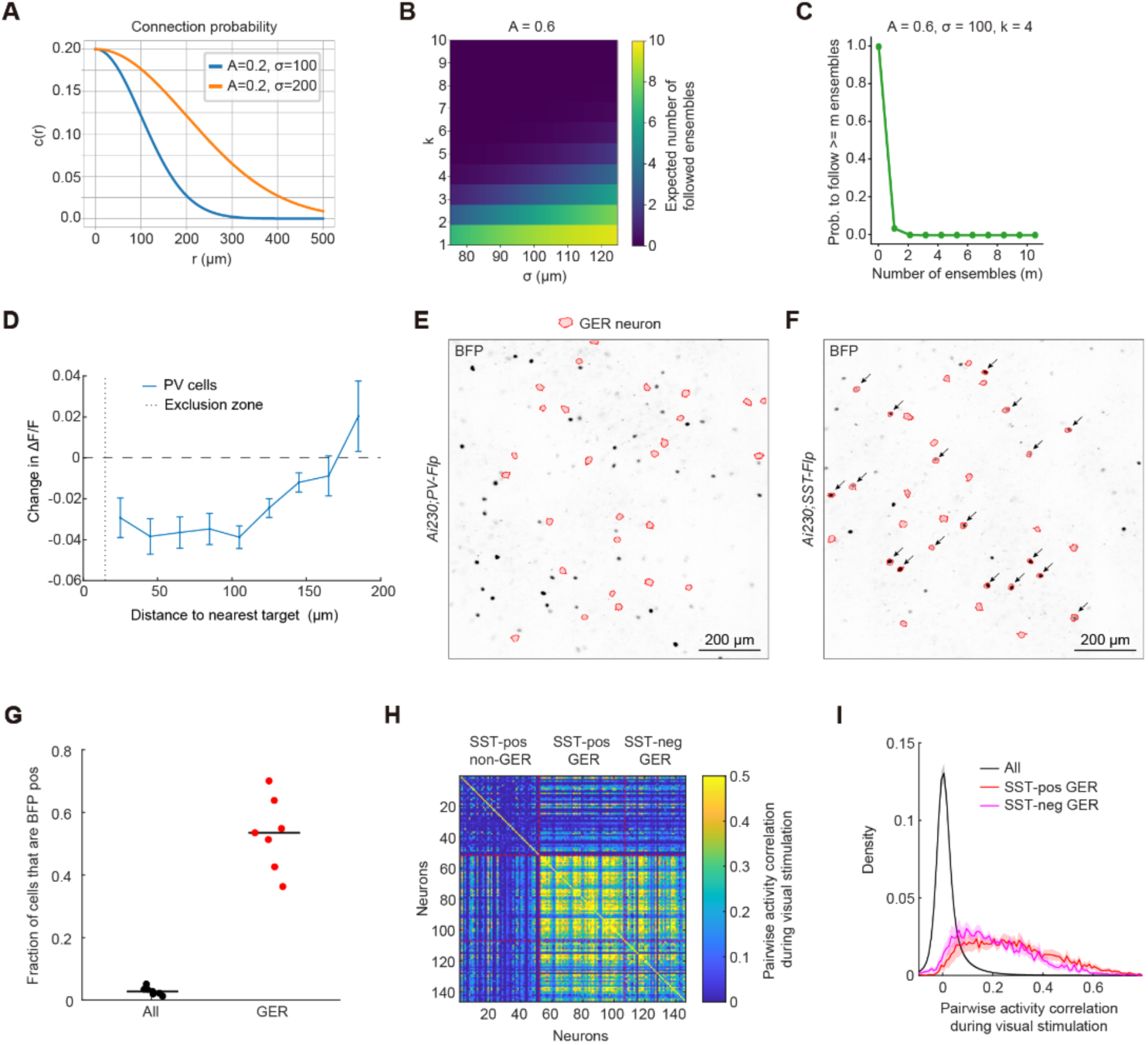
Related to Figure 6. **A**, Probability of connection, c(r), as a function of the distance (r in µm) between a pyramidal cell and a non-targeted neuron, for two different parameter sets of the Gaussian connectivity function: A = 0.2 and σ = 100 µm (blue), and A = 0.2 and σ = 200 µm (orange). **B**, Heatmap showing the expected number of ensembles (out of 10, each with 50 targets) that a non-targeted neuron will follow as a function of the activation threshold (k) and the spread of connectivity (σ in µm), for a peak connectivity (A) of 0.6. Despite increased peak connectivity, the likelihood of GER recruitment remained low (e.g., for k = 5, the expected number of followed ensembles was 0.4 out of 10), only increasing at very low k and large σ. **C**, Probability of following ≥m ensembles (out of 10) plotted as a function of m, for a peak connectivity (A) of 0.6, a spread of connectivity (σ) of 100 µm, and an activation threshold (k) of 4. **D**, Stimulation-evoked responses of PV cells as a function of distance to nearest stimulation target. Data represent mean ± SEM (n = 5 mice). **E**, Representative 2P image of the BFP signal in an example plane acquired in an *Ai230*;*PV-Flp* mouse intracranially injected with Flp-dependent AAV H2B-BFP (viral design in Figure 6F). Red masks show the Suite2p-detected ROIs of the GER neurons detected in this plane. No GER neuron was BFP-positive in this plane. **F**, Same as **E** but for a representative *Ai230*;*SST-Flp* mouse. Black arrows highlight BFP-positive GER neurons. **G**, Fraction of BFP-positive neurons per mouse, shown for the entire population (black) and GER neurons (red) in *Ai230*;*SST-Flp* mice intracranially injected with Flp-dependent AAV H2B-BFP (n = 7 mice). Black bars: median. Only neurons within the area encompassing all GER neurons were considered (**STAR Methods**). **H**, Correlation matrix of three groups of neurons during visual stimulation with drifting gratings for an example *Ai230*;*SST-Flp* mouse. Groups: SST neurons that were not classified as GER neurons (“SST-pos non-GER”), SST neurons that were classified as GER neurons (“SST-pos GER”), and GER neurons that were SST-negative (“SST-neg GER”). The visually-evoked activity of SST-positive and SST-negative GER neurons was highly correlated, indicative of a functionally coherent cell type and suggesting that the true proportion of SST+ GER neurons is likely to be even higher. **I**, Histograms of the pairwise activity correlation values during visual stimulation with drifting gratings for all neurons (black), GER neurons that were SST-positive (red), and GER neurons that were SST-negative (magenta) (mean ± SEM across 7 mice). The average pairwise correlation between SST-positive and SST-negative GER neurons was significantly larger than in the overall population (n = 7 mice, Wilcoxon signed-rank test, p = 0.016).

**Table S1.**
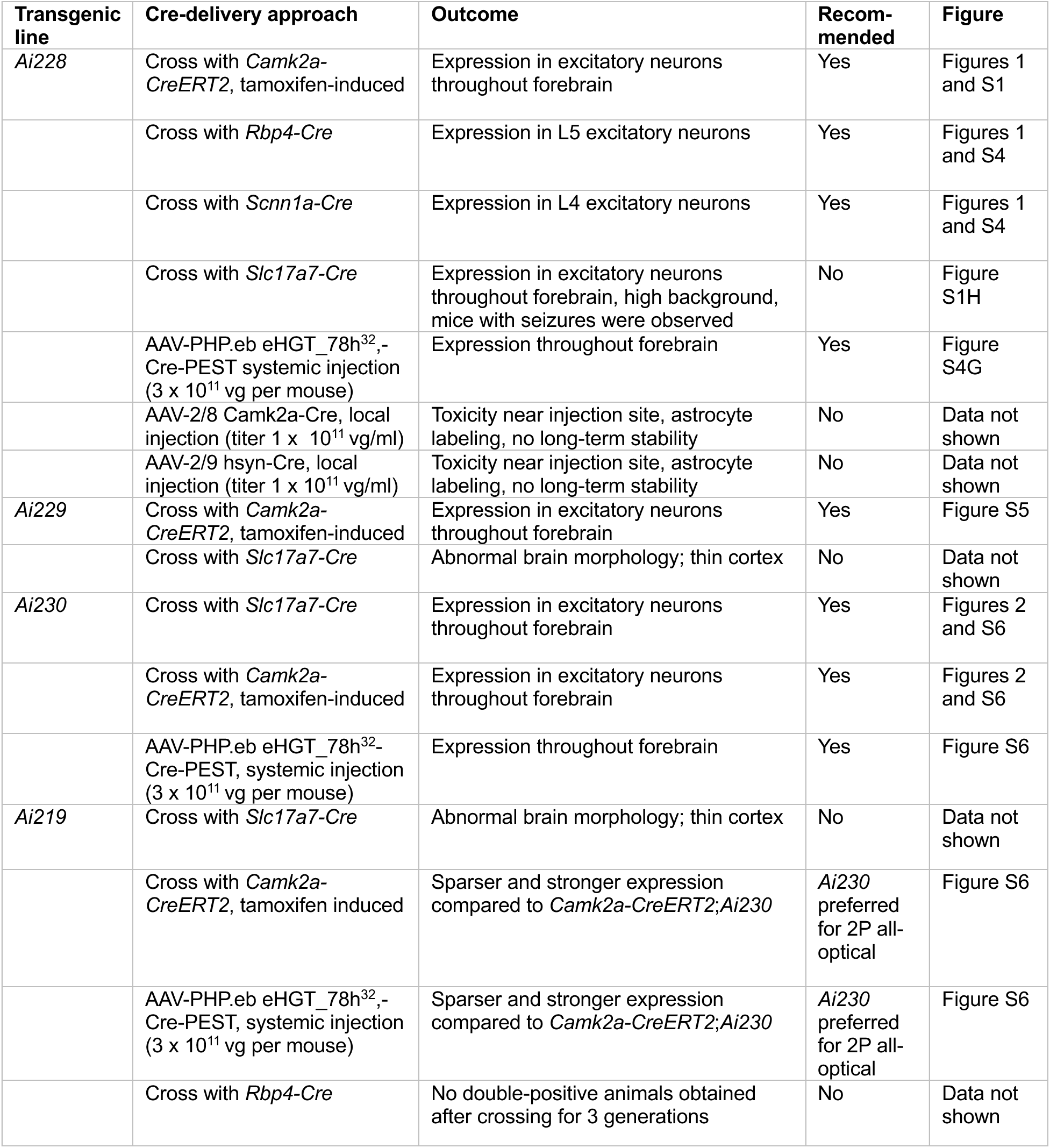
Summary of tested Cre-delivery strategies in *Ai228*, *Ai229*, *Ai230*, and *Ai219*.

## STAR METHODS

### Mice

Mouse strains generated in this study are *Ai228* (B6.Cg-Igs7<tm228(tetO-GCaMP6m,CAG-ChRmine*/oScarlet*,CAG-tTA2)Daigle>/J), *Ai229* (B6.Cg-Igs7<tm229(tetO-GCaMP6m,-ChRmine*,SYN1-tTA2)Daigle>/J), and *Ai219* (B6.Cg-Igs7<tm219(tetO-ChRmine*/oScarlet*,CAG-tTA2)Daigle>/J). Other mouse strains used in this study were *C57BL/6J* (JAX #000664), *Ai230* (JAX #37944), *Camk2a-CreERT2* (JAX #012362), *Slc17a7-Cre* (Slc17a7-IRES2-Cre-D, JAX #037512), *Scnn1a-Cre* (*Scnn1a-Tg1-Cre*, JAX #009111), *Rbp4-Cre* (*Tg(Rbp4-Cre)KL100Gsat*, No. 031125-UCD, MMRRC) (L. Luo), *PV-Flp* (*Pvalb-T2A-FlpO-D*, JAX #022730) and *SST-Flp* (*Sst-ires-Flp,* JAX #028579). Mice were group-housed in plastic cages with disposable bedding on a standard 12 h light cycle. All procedures were performed in accordance with protocols approved by the Stanford University Institutional Animal Care and Use Committee (IACUC) and guidelines of the National Institutes of Health.

### Generation of transgenic reporter mice

Ai228 (B6.Cg-Igs7<tm228(tetO-GCaMP6m,CAG-ChRmine*/oScarlet*,CAG-tTA2)Daigl>/J, construct TIT2L-GCaMP6m-ICL1-ChRmine-TS-oScarlet-Kv2.1-ER-ICL2-IRES2-tTA2) is a new TIGRE3.0 transgenic reporter^29^ for Cre-dependent expression of cytosolic GCaMP6m and soma-enriched ChRmine-oScarlet^13,28^. The modified TIGRE genomic locus in Ai228 contains (5’ to 3’): a Bxb1 attB site, a partial GFP sequence, two tandem copies of the chicken beta-globin HS4 insulator element, a Tet responsive 2 promoter (TRE2,comprised of seven repeats of TRE binding sites and a minimal CMV promoter based on Clontech’s pTRE2-hyg vector), a LoxP-flanked stop cassette, a truncated GCaMP6m coding sequence, a woodchuck post-transcriptional regulatory element (WPRE), and a polyA signal. It also includes two tandem copies of the chicken beta-globin HS4 insulator element, a CAG promoter (consisting of the CMV enhancer fused to the chicken beta-actin promoter), a Lox2272-flanked stop cassette, and the coding sequence for ChRmine fused to a membrane trafficking signal (TS), oScarlet, Kv2.1 and an ER export signal, followed by a WPRE and polyA signal. Further, the locus contains two tandem copies of the chicken beta-globin HS4 insulator element, a CAG promoter, a LoxN-flanked stop cassette, an IRES sequence, tetracycline-transactivator 2 (tTA2), a WPRE, and a polyA signal. Finally, it includes an AttB site, a PGK promoter driving one domain of the hygromycin resistance gene, and a Bxb1 attB site. The locus was generated via recombinase-mediated cassette exchange into a Bxb1 integrase docking site integrated into the TIGRE locus in mouse embryonic stem (ES) cells^79^.

Ai229 (B6.Cg-Igs7<tm229(tetO-GCaMP6m,-ChRmine*,SYN1-tTA2)Daigl>/J, construct TIT2L-GCaMP6m-IRES2-ChRmine-TS-Kv2.1-HA-ISL-tTA2) is a new TIGRE2.0 transgenic reporter for Cre-dependent expression of cytosolic GCaMP6m and soma-enriched ChRmine. The modified TIGRE genomic locus in Ai229 contains (5’ to 3’): a Frt3 site, two tandem copies of the chicken beta-globin HS4 insulator element, TRE2, a loxP-flanked stop cassette, a truncated GCaMP6m coding sequence, an IRES2 sequence, the coding sequence for ChRmine fused to TS, oScarlet, and Kv2.1, followed by WPRE and a bGH pA sequence. Additionally, the locus includes two tandem copies of the chicken beta-globin HS4 insulator element, a fragment of the human synapsin 1 gene promoter^80^, a lox2272-flanked stop cassette, tTA2, a WPRE, and a bGH pA sequence. Finally, it contains a PGK promoter driving one domain of the hygromycin resistance gene, an mRNA splice donor sequence, and a Frt5 site. The locus was generated by recombinase-mediated cassette exchange into a previously established docking site within the TIGRE locus^29^ in mouse ES cells.

Ai219 (B6.Cg-Igs7<tm219(tetO-ChRmine*/oScarlet*,CAG-tTA2)Daigl>/J, construct TIT2L-ChRmine-TS-oScarlet-Kv2.1-ER-ICL-IRES2-tTA2) is a new TIGRE2.0 transgenic reporter for Cre-dependent expression of soma-enriched ChRmine-oScarlet. The modified TIGRE genomic locus in Ai219 contains (5’ to 3’): a Frt3 site, two tandem copies of the chicken beta-globin HS4 insulator element, TRE2, a loxP-flanked stop cassette, the coding sequence for ChRmine fused to TS, oScarlet, Kv2.1 and an ER export signal, followed by a WPRE and a bGH pA sequence. Additionally, the locus includes two tandem copies of the chicken beta-globin HS4 insulator element, a CAG promoter, a lox2272-flanked stop cassette, an IRES sequence, tTA2, a WPRE, and a bGH pA sequence. Finally, it contains a PGK promoter driving one domain of the hygromycin resistance gene, an mRNA splice donor sequence, and a Frt5 site. As for Ai229, generation of the locus involved recombinase-mediated cassette exchange into a previously established docking site within the TIGRE locus^29^ in mouse ES cells.

Generation of all new lines involved injecting correctly targeted ES cell clones into fertilized blastocysts, which produced high-percentage chimeras. These chimeras were subsequently bred with C57BL/6J mice to achieve germline transmission. The resulting heterozygous TIGRE reporter mice were first maintained in a C57BL/6J congenic background, then intercrossed to generate homozygous mice. For Cre induction, TIGRE reporter mice (*Ai228, Ai219, Ai230,* or *Ai229*), either homozygous or heterozygous, were crossed with homozygous or heterozygous Cre driver lines (*Camk2a-CreERT, Slc17a7-Cre, Scnn1a-Cre, Rbp4-Cre*). Alternatively, heterozygous TIGRE reporter mice were injected with AAV (**Table S1**).

### Viral constructs

AAV-2/8 hSyn-GCaMP8s-3xGGGS-L10a-ARE was generated using standard cloning techniques. This construct uses the L10 sequence^36^ to restrict GCaMP8s expression to the soma via ribosome-tagging. We made two changes in the original design to optimize the expression: First, we added a 3xGGGS linker between the GCaMP and L10a (similar to reference^52^). Second, we added a 3’UTR sequence (A2RE^81^, an 11 nucleotides long RNA trafficking signal binding the heterogeneous ribonucleoprotein A2), which stabilizes mRNA and therefore increases expression levels. AAV-2/8-Ef1a-fDIO H2B-BFP was generated using standard cloning techniques. AAV-PHP.eb eHGT_78h-Cre-PEST was published previously^34^; the construct is available on Addgene (#231791) and prepackaged virus can be purchased from the Stanford GVVC.

### Retro-orbital injections

For systemic delivery via retro-orbital injections, AAV-PHP.eB-eHGT_78h-Cre-PEST was prepared to a final concentration of 6 x 10^11^ vg/mouse in a total volume of 40 µL ice-cold PBS. The viral solution was front-loaded into a BD Micro-Fine insulin syringe (0.5 mL, 30G). Mice (4-7 weeks old) were anesthetized with isoflurane for the procedure. The viral solution was administered into the retro-orbital sinus, with the needle inserted from the nasal side. Subsequent craniotomies were performed 3-6 weeks after retro-orbital injections.

### Tamoxifen injections

Tamoxifen solution (20 mg/mL) was prepared by dissolving 100 mg of tamoxifen powder (Sigma, T5648) in 5 mL of pre-heated corn oil in a scintillation vial. The vial was tightly sealed, vortexed for at least 30 s, and incubated on a rocker at 37°C overnight to aid dissolution. The following day, the solution was drawn into a 5 mL syringe without a needle. An 18G needle was then attached and the solution was pushed through the needle to disperse any undissolved clumps. This process was repeated up to five times until all clumps were dissolved. To protect from light, the vial was wrapped in aluminum foil. The prepared tamoxifen solution was stored at 4°C and used within one month. Tamoxifen solution was administered via intraperitoneal (i.p.) injection at a concentration of 80 mg/kg once every 48 hours for three days to double-heterozygous *Camk2a-CreERT2*;ChRmine-reporter mice (*Ai228*, *Ai229*, *Ai219,* or *Ai230*). Injections were performed either one week before or one week after craniotomy. Mice were imaged after a waiting period of at least one week after the final tamoxifen injection.

### Surgeries

Female and male mice (8-12 weeks old) were used. Mice received dexamethasone (s.c. 2 mg/kg) 2-3 hours before surgery to reduce inflammation. For preparations with intracranial injections, 5 μL of viral solution was prepared. Viral solution consisted of AAV-2/8 hSyn-GCaMP8s-3xGGGS-L10a-ARE (3 x 10^12^ vg/ml) for labeling with ribosome-tagged GCaMP8s, or AAV-2/8 hSyn-GCaMP8s-3xGGGS-L10a-ARE (3 x 10^12^ vg/ml) gently mixed with AAV-2/8 Ef1a-fDIO H2B-BFP (1 x 10^11^ vg/ml), both in ice-cold PBS, for additional Flp-dependent BFP labeling. Viral solution was front-loaded into a glass pipette (15-20 µm opening, beveled tip) using a Nanoinject III injector (Drummond Scientific, Catalog No. 13-681-460). Mice were then anesthetized using a mixture of Ketamine (100 mg/kg) and Xylazine (10 mg/kg), supplemented with 0.5-1% isoflurane. The skull was exposed, and a 5 mm craniotomy was performed above visual cortex using a dental drill, ensuring the dura remained fully intact. During surgeries with intracranial injections, 1 μL viral solution was injected at a speed of 1 nl/s at 400 µm below the brain surface in the primary visual cortex. The pipette was left in place for 5-10 min after injection and then slowly retracted. The craniotomy was then closed with a 5 mm glass coverslip. A metal head bar was mounted using dental cement (Metabond). 30 min before the end of surgery, Buprenorphine sustained release (SR) (0.4 mg/kg) was injected subcutaneously. Craniotomies were inspected under the stereomicroscope 1-2 weeks after surgery. In the case of tissue growth, the window was re-opened under anesthesia. Growing tissue was carefully removed using fine forceps before a new glass window was put in place. 2P imaging was performed >2 weeks after surgery for fully-transgenic preparations, and >4 weeks after surgery for preparations with intracranial injections.

### All-optical two-photon holographic experiments

#### Overview microscopy system

The microscope incorporated a custom-built stimulation pathway integrated with a commercial two-photon (2P) microscope (Bruker Ultima 2P plus, controlled by PrairieView software). This system was similar to the multiSLM setup described previously^13^, with the key differences being the increased imaging field-of-view (FOV, maximum FOV 1.4 x 1.4 mm² using a Nikon CFI75 LWD 16x objective) and use of an electro-tunable lens for axial scanning instead of the piezo element. Moreover, a motorized nosepiece was used to optimize the objective tilt.

#### Imaging

Imaging was performed using a tunable laser (Coherent Discovery with TPC). The following wavelengths were used: 920 nm for GCaMP imaging (green channel), 980 nm for oScarlet imaging (red channel) or simultaneous GCaMP and oScarlet imaging (green and red channels), and 870 nm for BFP imaging (blue channel). Imaging power (measured below the objective during scanning) was adjusted for each imaging plane; the ranges used varied depending on the mouse line and cortical layer: In *Ai228* mice, 20-30 mW was used in L2/3, and 30-40 mW in L4 and L5; In *Ai229* mice, 30-40 mW was used in L2/3; In *Ai230* mice with pan-neuronal ribosome-tagged GCaMP8s, 30-55 mW was used in L2/3. Imaging was performed using a 16x/0.8 NA objective (Nikon CFI75 LWD) immersed in diluted ultrasound gel. Unless otherwise stated, the FOV was 1 x 1 mm². In L2/3 experiments, the volume consisted of 3 or 5 imaging planes. The first plane was positioned approximately 150 µm below the surface, and subsequent planes were spaced 30 µm apart. Images were acquired at a resolution of 1024 x 1024 pixels, at 3 Hz for 5-plane imaging and 5 Hz for 3-plane imaging.

#### Holographic stimulation

Stimulation light was generated by a fixed-wavelength (1035 nm) femtosecond laser (Coherent Monaco 1035-80-60) operating at a 2 MHz pulse repetition rate. Holograms were generated using a spatial light modulator (SLM, Boulder Nonlinear Systems / Meadowlark Optics macroSLM, 1536 x 1536 pixels with temperature control). Holograms were computed from 3D target coordinates using an algorithm described previously^13^. The hologram was scanned in a spiral (5 rotations, 5 µm diameter) across targeted neuron bodies using a pair of galvo mirrors for 2 ms. To address the systematic drop-off of SLM diffraction efficiency across the FOV^49^, the galvo mirrors would also pan the gaze of the hologram to be centered within the target ensemble (XY centers shifted by values up to ± 120 µm), ensuring efficient photo-stimulation across the field. In a subset of experiments (data shown in **Figures 3, 4, 5 and 6**), the target locations were split diagonally into two groups, each stimulated with a separate hologram. Both holograms were panned to their respective target centers and temporally spaced 2 ms apart to provide sufficient time to cycle through the respective phase masks; the entire ensemble was stimulated within 6 ms (“near-simultaneous”). Stimulation parameters (duration, frequency, power) varied across experiments (see text for details). The inter-trial interval (ITI) ranged from 4.5 to 8 s.

For power scaling of the hologram across different depths, an amplitude term weighted by the function 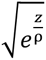 with scattering length ρ = 156-250 µm was added to each target in the hologram. The time-averaged stimulation power per neuron was computed by

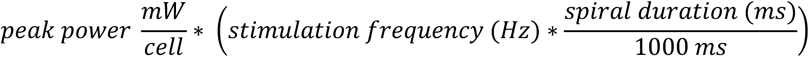

Optical aberrations of the optogenetic beam were corrected by scanning the optogenetic galvo pair and forming an image using a bulk fluorescent slide. Sequentially applying varying weights of Zernike polynomials (OSA/ANSI single-indexed modes 3-20) to the SLM phase mask resulted in performance curves used to select values which maximize the resulting image intensity (i.e. minimize optical aberration), as in reference^82^.

#### Alignment of stimulation and imaging paths

To ensure precise alignment before each stimulation experiment, a predefined pattern of holes was burnt into a fluorescent slide using the stimulation laser. The centers of these holes were then manually identified in the 2P image, and the coordinates of the holes were compared to the intended location of the calibration pattern. This allowed us to identify and correct lateral shifts between the imaging and stimulation paths (typically 1-3 pixels).

#### Online motion correction

At the start of each experiment series, a reference image stack (25 planes, Δz = 1 µm, spanning ±12 µm around the center imaging plane, averaged over 100 frames) was acquired. A custom MATLAB script, interfacing with Prairie View, performed online motion correction in 3D by registering live center-plane images during the experiment to the reference stack using a cross-correlation-based algorithm. This approach was also used to align the FOV across days, after moving the sample in the approximate location using blood vessels as landmarks.

#### Cell selection for holographic stimulation

Cells were selected in two possible ways: (1) online, manual selection while the mouse remained in the setup (experiments shown in **Figures 1, S2, S4 and S5**), or (2) offline selection while the mouse was removed from the setup (experiments shown in remaining figures).

For online selection, GCaMP+ neurons were manually selected from a 128-frame average image of the green channel using the ROI selection tool of PrairieView (‘BOT tool’). Using a custom MATLAB script, holographic target coordinates were then calculated from the centroids of the marked neurons.

Offline selection involved acquiring (1) time-series data (green channel acquired at 920 nm, usually during visual stimulation), which was processed as described in “Analysis of two-photon imaging data” (2) a brief time-series recording of the green and red signals (500 frames per plane, 980 nm simultaneous imaging), which was registered using Suite2p^76^ (non-rigid), and (3) a 128-frame average image (green channel, each plane). From the motion-corrected red channel images, oScarlet+ neurons were identified using Cellpose^78^ with a custom model, followed by manual curation. The oScarlet+ neurons were then registered to Suite2p-detected neurons of the time-series data (GCaMP-ROIs) in each plane: First, the 980 nm time-series data was aligned to the 920 nm time-series data by registering the motion-corrected green channel image of the 980 nm time-series to the motion-corrected image of the 920 nm time-series data using cross-correlation. Subsequently, the closest GCaMP-ROI to each oScarlet+ neuron was found. If no GCaMP-ROI was found within 5 pixels (4.88 µm), no assignment was made. The oScarlet+ GCaMP-ROIs constituted the pool of potential stimulation targets. From this pool, targets were chosen randomly or based on their functional properties (see section “Selection of stimulation ensembles”). Since the motion-corrected image of the 920 nm time-series data was not always aligned to the reference stack that we used for online motion correction (see section “Online motion correction“), in a last step, X and Y shifts between the 128-frame average (green channel) and the motion-corrected image of the 920 nm time-series data were computed per plane. The centroids of the shift-corrected GCaMP-ROIs were then used to compute the holographic target coordinates.

#### BFP imaging and cell detection

A brief time-series recording of the green and blue signals (500 frames per plane, 870 nm simultaneous imaging) was acquired. From motion-corrected (Suite2p, non-rigid) blue channel images, BFP+ neurons were identified using Cellpose (‘cyto2’ model) and then manually curated. The BFP+ neurons were subsequently assigned to Suite2p-detected GCaMP-ROIs from the time-series data in each plane. This involved aligning the 870 nm time-series data to the 920 nm time-series data, then finding the closest GCaMP-ROI to each BFP+ neuron. If no GCaMP-ROI was found within 10 pixels (9.8 µm), no assignment was made.

#### In vivo two-photon (2P) images of brain preparations

To visualize *in vivo* expression of GCaMP, ChRmine-oScarlet, and BFP, 500 frames 2P time-series in each channel were acquired as described above. Frames were registered using Suite2p (non-rigid). Displayed images represent the average of the registered frames. The motion-corrected frames of different channels were registered using cross-correlation. Photographs of cranial windows were taken using a cell phone camera through a stereoscope.

#### Cross-stimulation measurements and analysis

To assess unintended activation of ChRmine by the imaging laser, 16 frames at 5 Hz in a 3-plane volume (30 µm spacing, 1 x 1 mm² FOV) were acquired using varying imaging laser power levels (**Figure S3**). Power levels were randomized across 10 trials per level, with a 10 s inter-trial interval. Suite2p was used for registration, neuron detection, and extraction of fluorescence traces. The F_AVG,j_ time series was estimated for each power level j by averaging fluorescence traces across trials and neurons. This series was then normalized by min(F_AVG,j_) to compute ΔF_AVG,j_. The cross-stimulation effect for each power level j was then estimated as the difference between the last and first values of the ΔF_AVG,j_ time series. For trial-specific analysis, trialF_i,j_ for each power level j and each trial i was computed by averaging fluorescence traces from all neurons for each trial. Subsequently, trialΔF_i,j_ was derived by normalizing trialF_i,j_ by min(F_AVG,j_).

### Visual stimulation during two-photon imaging

Head-fixed mice, free to run on a foam cylinder, viewed visual stimuli presented on a monitor (HP VH240a) positioned 10 cm from the eye. Stimuli were controlled by custom-written software in python.

#### Drifting gratings

Grating stimuli were circular patches (35° diameter, 80% contrast) of sinusoidal gratings drifting in 8 orientations (45° spacing, 0.03 cpd spatial frequency, 1 Hz temporal frequency). Patches were presented for 4 s on a gray background, interleaved with a 4 s gray screen. Each orientation was presented 15 times in random order.

#### Natural Scenes

The stimulus consisted of a sequence of 118 grayscale natural images (Allen Brain Observatory library^40^). Each image was presented for 0.75 s with a 1.75s gray period in between. The same sequence was repeated 10 times. In one out of 13 mice, a sequence of 236 images (the original 118 and their horizontal flips) was presented 5 times.

#### Retinotopic mapping

Circular patches (15° diameter, 100% contrast) displaying sinusoidal gratings were used (0.03 cpd spatial frequency, 2 Hz temporal frequency). During each stimulation trial, gratings of four distinct orientations (0°, 45°, 90°, 135°) were presented, with each orientation displayed for 0.5 s, for a total stimulus duration of 2 s. These patches were presented at nine locations, with their centers arranged in a 3 x 3 grid tiling the monitor, on a gray background. Each location was shown 12 times in random order, interleaved with a 4 s gray screen.

### Analysis of two-photon imaging data

#### ΔF/F

Time-series data were analyzed using Suite2p^76^ for non-rigid registration, automated neuron segmentation, and extraction of calcium fluorescence traces. To track neurons across multiple recording sessions, data were concatenated and analyzed as a single long recording in Suite2p. Subsequent analysis was performed in MATLAB. Neuropil contamination was reduced by estimating and subtracting neuropil signals for each ROI, following the method described in this paper^11^. Neuropil subtraction coefficients were determined using robust regression (MATLAB’s ‘robustfit’ function with ‘bisquare’ weighting) between the downsampled (10x) ROI signal and the downsampled (10x) neuropil signal. Coefficients were constrained between 0.5 and 1; if regression failed to converge, the median of well-fit coefficients was used. The resulting neuropil-corrected traces were then re-baselined to match the baseline of the original, uncorrected traces. To correct for slow baseline drift, a running baseline (20 s sliding window) was subtracted from the neuropil-corrected calcium traces^76^. The resulting traces were then normalized to their median value to obtain ΔF/F values.

#### Threshold for exclusion

For quality control, neurons with unusually high ΔF/F values were excluded from analysis. We first determined the 99.99 percentile of the distribution of a large subset of the data (T = 9.98 ΔF/F). Cells were excluded if more than 0.5% of datapoints per imaging session were above this threshold.

#### Analysis of visual responses

ΔF/F responses to grating and natural scene stimuli were quantified as the difference between the average ΔF/F during post-stimulation (grating stimuli: 0-4 s; natural scenes: 0-1.2s) and pre-stimulus baseline (grating stimuli: -2 to 0 s; natural scenes: -0.5 to 0s) for each trial and then averaged across trials.

#### Orientation-selectivit

An orientation selectivity index (OSI) was calculated using the formula:

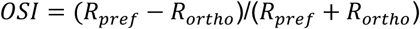

where *R_pref_* and *R_orhto_* denote the response amplitudes for the preferred and orthogonal orientations, respectively. Non-responsive neurons to grating stimuli (max. ΔF/F response <0.5 across orientations) were excluded when quantifying the fraction of orientation-selective neurons (**Figure 5C**).

#### Sparseness of responses to natural scenes

Sparseness (S) of responses to natural scenes was calculated according to reference^41^ as:

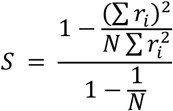

where N is the number of scene stimuli and r_i_ is the neuron’s trial-averaged response to the i^th^ stimulus. Values of *S* near 0 indicate a dense code, and values near 1 indicate a sparse code. Non-responsive neurons to natural scene stimuli (max. ΔF/F response <0.5 across scenes) were excluded when quantifying the sparseness (**Figure 5D**).

#### Pairwise activity correlation

Pairwise activity correlation was computed from ΔF/F time-series data during visual stimulation and averaged. For spontaneous activity correlation, the final 3 s of the 8 s inter-stimulus interval from holographic stimulation sessions were concatenated per trial, and pairwise correlation was calculated using these concatenated time-series.

#### Retinotopic maps

Responses to each of 9 grating locations were computed as the difference between the average ΔF/F during post-stimulation (0-1 s) and pre-stimulus baseline (-1 to 0 s) for each trial and then averaged across trials. The preferred stimulus location for each neuron along either the horizontal or vertical axis was determined as the location eliciting the largest average ΔF/F response. Neurons with a maximal response less than 0.3 ΔF/F across all tested locations were excluded. Borders between V1 and HVA were manually drawn on horizontal retinotopic maps guided by the reversal in the representation of the vertical meridian.

#### Selection of stimulation ensembles

Scene-selective ensembles were constructed using hierarchical clustering of z-scored, trial-averaged single-neuron responses natural scenes. Responses were represented as a *N* × (*n_scenes_* · *t*) matrix, where *N* is the number of neurons and *t* represents the number of imaging frames from stimulus onset (*t* = 5 was used, with 4 pre-stimulus frames for baseline correction). Hierarchical agglomerative clustering was performed using the ‘scipy.cluster.hierarchy’ library, with average linkage and one minus the Pearson correlation as the distance metric. The neuron-neuron correlation matrix, ordered by the dendrogram leaves, was inspected for block structure, which ensured that clustering merges did not purely occur at large linkage distances, but instead over a broad range. Clusters were extracted by thresholding the dendrogram at linearly spaced linkage distances between 0.3 and 0.9 (bounds varied slightly across mice to tune the number of candidate clusters). Clusters with 170 to 500 neurons were retained (bounds varied across mice to yield approximately 10 to 12 candidate clusters). A quality metric, defined as the average pairwise correlation of its neurons, excluding self-correlations, was computed for each cluster. Clusters were then sorted by quality score and subsets of higher-quality clusters were discarded. Clusters were further filtered to contain at least *k* oScarlet+ neurons (*k* ranged from 30-50 across mice, depending on the target ensemble size per mouse). Finally, after visually inspecting the trial-averaged and single-trial activity per cluster, two to four ensembles were selected per experiment, prioritizing clusters with consistent scene-specific responses across trials.

To construct orientation-selective ensembles, selectivity metrics (response at each orientation, preferred orientation, and OSI) were computed using *t* = 5 frames post-stimulus onset and 4 pre-stimulus frames for baseline correction. In 9 of 10 mice of the V1 dataset and in 2 of 3 mice of the V1-LM dataset, responses were averaged across both directions for each orientation (collapsing the eight grating directions into four distinct orientations), resulting in orientation-but not direction-selective ensembles. In the remaining mice, responses were computed separately for all eight grating directions, resulting in direction-selective ensembles. Neurons were discarded if they did not meet two criteria: (1) a minimum OSI (between 0.13 and 0.2) and (2) a minimum response amplitude at the preferred orientation (0.25). These thresholds were adjusted per mouse to prevent disproportionate neuron loss based on the distribution of OSI and response amplitudes. Neurons meeting these thresholds and determined to be oScarlet+ neurons were grouped by preferred orientation. Within each group, each neuron’s stimulus-averaged, z-scored trace was correlated with the group’s mean trace. Neurons within each group were sorted by this correlation, and stimulus-averaged traces were visualized. The top 30-50 neurons per orientation (depending on the target ensemble size per mouse) were selected as an ensemble.

Ensembles were mutually exclusive. Scene-selective ensembles were constructed first, and their neurons were then excluded from the selection of orientation-selective ensembles. Size-matched random ensembles were constructed using the remaining oScarlet+ neurons, excluding those in the functional ensembles.

#### Artifact removal

The holographic stimulation produces a stimulation artifact during each 2 ms pulse. The imaging frequency (3 Hz or 5 Hz) and stimulation frequency were selected such that stimulation artifacts did not overlap in adjacent frames (i.e. consecutive frames in time) within each imaging plane. Prior to registration and ROI extraction, stimulation artifacts were removed by replacing the pixel rows containing the stimulation artifact with the average values from corresponding rows in the preceding and following frames. The artifact-free imaging data were then processed as described above.

#### Assignment of targeted neurons

In each plane, stimulation targets were assigned as the GCaMP-ROIs with centroids closest to the designated target coordinates. If no GCaMP-ROI was located within a 5-pixel radius of a given target coordinate (for example, due to a cell’s inactivity and subsequent failure of detection by Suite2p), no targeted neuron was assigned to this target coordinate.

#### Quantification of stimulation success

Stimulation-evoked ΔF/F responses were quantified as the difference between the average ΔF/F during post-stimulation (0.2-2 s) and pre-stimulus baseline (-1.5 to 0 s). Response reliability of a stimulation target was quantified as the mean pairwise correlation coefficient of the stimulation-evoked ΔF/F activity trace (computed from stimulation onset to 3 s after onset) across all trials. A targeted neuron was deemed responsive if this value exceeded 0.2.

#### Fluorescence and ΔF images

Fluorescence images were generated by averaging registered frames (post-stimulation period: 0.2-2 s) across trials. ΔF images, derived from the difference between post- and pre-stimulation fluorescence images (pre-stimulation period: -1.5 to 0 s), were averaged across trials and smoothed using a 2D Gaussian filter (MATLAB ‘imgaussfilt’, σ = 0.75).

#### Exclusion zone

Distances between neurons were computed as the distance between the GCaMP ROI centers in the 3D imaging volume. The non-targeted population was defined as all ROIs within the imaged volume, excluding the stimulation targets and neurons at high risk of direct photostimulation. Those high-risk neurons were defined as neurons within a 30 µm diameter cylinder centered on the X, Y location of each target, extending through all imaging planes (“exclusion zone”). An extra conservative exclusion zone was chosen for the classifier analysis (**Figure 3**) to further minimize the risk of including directly photostimulated neurons. Here, we excluded neurons within a 60 µm diameter cylinder extending through all planes.

#### Classifier analysis

Ensemble identity was decoded from the stimulation-evoked responses of non-targeted neurons using a support-vector machine (SVM) classifier. The classifier was trained and evaluated on single-trial responses (n = 20-25 trials per ensemble) using 5-fold cross-validation. For each fold, 80% of the trials served as the training set and 20% as the test set. Model training and prediction were performed using the MATLAB functions ‘fitcecoc’ and ‘kfoldPredict’, respectively. Chance-level decoding performance was established by shuffling trial labels 10 times and averaging the resulting classifier performance across these repetitions. Classifier performance was visualized using a confusion matrix generated with the MATLAB function ‘confusionmat’. For the sliding window analysis, training and evaluation were performed as described above, except that single-trial responses were replaced with single-frame activity, and this analysis was repeated for each frame from -1 s to 3 s relative to the stimulation onset.

#### Evoked ΔF/F as a function of distance

or analyzing stimulation-evoked responses relative to the nearest target, evoked ΔF/F and the distance to the nearest targeted neuron were calculated for all non-targeted neurons (excluding neurons in the exclusion zone) for each stimulation ensemble. For analysis involving GER neurons (**Figures 4D and 6N**), these values were computed for each ensemble *X* using GER neurons identified in the remaining ensembles (excluding *X*). Data was then binned by distance (bin width 10 or 20 µm) and averaged per bin for each ensemble, giving an estimate of the response vs distance curve per ensemble.

#### Follower analysis

To identify positive and negative followers within the non-targeted population for each stimulated ensemble, a 99% confidence interval was computed for the mean stimulation-evoked ΔF/F response of each neuron across trials using a *t*-distribution. The *t*-value was determined based on a significance level of p = 0.01 and degrees of freedom (number of trials – 1). The confidence interval was then computed as the mean ± (*t*-value * SEM). Based on these confidence intervals, positive followers were defined as neurons from the non-targeted population for which the lower bound of the confidence interval was greater than zero, indicating a statistically significant increase in stimulation-evoked ΔF/F. Conversely, negative followers were defined as neurons for which the upper bound of the confidence interval was less than zero, indicating a statistically significant decrease in stimulation-evoked ΔF/F.

GER neurons were defined as positive followers recruited by ≥4 ensembles. The chance level for follower detection (**Figure S9B**) was established by analyzing surrogate data generated from randomly sampled time points in the original recordings (artificial stimulation onsets). This process was repeated 1000 times to generate multiple surrogate datasets for reliable chance-level estimation. The fraction of positive followers recruited by at least *N* ensembles (**Figure 4C**) was computed by counting for each neuron within the non-targeted population the number of times that it was considered a positive follower across the different stimulated ensembles. The resulting counts were normalized by the total number of positive followers across all ensembles. The expected rate of follower overlap across ensembles due to chance (shuffle control in **Figure 4C**) was estimated by randomly reassigning neuron identities in the non-targeted population for each ensemble while maintaining the original number of positive followers per ensemble. This shuffling was repeated 100 times, and the same counting and normalization procedure was applied to estimate the expected follower rate.

#### Linear mixed-effects models

Linear mixed-effects models (used to compare the classifier performance and follower counts across ensemble categories) were implemented with the MATLAB function ‘fitlme’. The models aimed to predict either classifier performance (**Figure S8C**) or follower count (**Figures S9C and S9D**). The fixed effects included ensemble category, the number of targets within each ensemble, and the average evoked ΔF/F response in the targeted neurons. To account for potential variability between individual mice, a random intercept for mouse identity was included in the model. For this analysis, OS-selective and scene-selective ensembles were combined and treated as functional ensembles.

#### GER enrichment across depth

To quantify the spatial distribution of GER neurons across imaging planes (**Figure 5G**), we calculated an enrichment index. For each plane, we first determined the relative proportion of GER neurons in that plane by dividing the number of GER neurons in that plane by the total number of GER neurons across all planes. We performed a similar calculation for all neurons. The enrichment index for each plane was then calculated as the difference between these two proportions, normalized by the relative proportion of all ROIs for that plane. This metric highlights planes where GER neurons are overrepresented relative to the total population of recorded neurons.

#### Analysis of BFP+ neurons

Only BFP+ neurons outside the exclusion zone were included in the analysis. To account for differences in spatial target distribution across mice, BFP+ neurons outside of the area encompassing all GER neurons were excluded from analysis. This area was defined for each mouse as the convex polygon encompassing the projected locations of all GER neurons (across all imaging planes) onto one plane.

### Computational model

A probabilistic network model was used to estimate the likelihood of a non-targeted neuron getting activated by (“following”) the targeted stimulation of an ensemble of randomly chosen pyramidal cells. The model assumed random, purely spatial separation-dependent connectivity between targeted pyramidal cells and non-targeted neurons. The probability of a synaptic connection between a pyramidal cell and a non-targeted neuron as a function of their separation, *r*, was modeled using a Gaussian function: *p*(*r*) = *Ae*^-*r*2/(2σ2)^, where *A* is the peak connection probability and *σ* is the spread of connectivity. A three-parameter model of a non-targeted neuron was defined using parameters *A* and *σ* (as described above), and *k*, which represents the minimum number of connected pyramidal cells that must be stimulated to activate the non-targeted neuron. An ensemble was defined as a set of *T* targeted pyramidal cells, with the *T* cells chosen uniformly and randomly within the FOV. We considered a total of *M* sampled ensembles, each containing *T* cells.

The objective was then to derive the probability distribution of the number of ensembles *m* followed by a non-targeted neuron, as a function of the model’s parameters, that is *p*(*m*; *A*, *σ*, *k*, *M*, *T*) for *m* ≤ *M*, and particularly the expected value of this distribution, 𝔼(*m*; *A*, *σ*, *k*, *M*, *T*). For given choices of *A*, *σ*, and *k*, where *M* and *T* were chosen according to experimental settings (*M* = 10 ensembles, *T* = 50 targets) the likelihood of GER neurons could be quantified as follows:

A non-targeted neuron was placed in the center of a square FOV with side length *L*. Targeted pyramidal cells were distributed randomly and uniformly within this FOV, with a density *ρ*. First, consider a single pyramidal cell, so that *ρ* = 1/*L*^2^. The probability of this targeted cell being located within a disk of radius *r* and width *dr* is *ρ* · 2*πrdr*. Multiplying this term by the connection probability at that radius gives the probability that the targeted cell is connected to the non-targeted neuron at the origin:

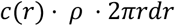

Therefore, the overall probability of a connection between a randomly placed targeted cell and the non-targeted neuron can be obtained by integrating this probability over the field-of-view:

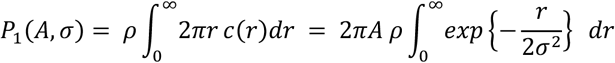

The upper integration bound can be taken to be ∞ if σ is sufficiently small relative to *L*. For an ensemble of *T* cells, we can now compute the probability, *P_f_*, that at least *k* cells are connected to the non-targeted neuron. This is equivalent to one minus the probability that the number of successes with *T* trials is less than *k*,

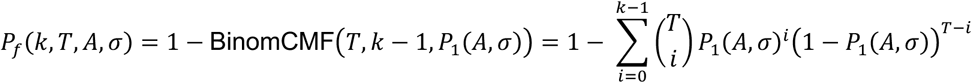

where BinomCMF(*N, n, p*) is the cumulative mass function for a binomial distribution with *N* trials, probability of success *p*, and at most *n* successes.

Finally, the probability that the non-targeted neuron follows *m* out of *M* ensembles is given by the binomial probability of *m* successes in *M* independent trials, where the probability of success (following in ensemble) is *P_f_*, is:

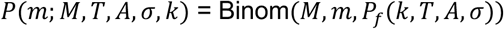

The expected number of ensembles followed, out of the total *M* ensembles, is then:

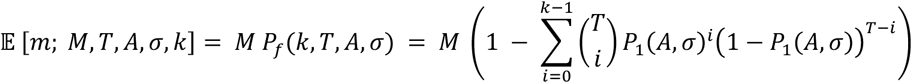

### Histology and confocal imaging

#### Perfusion

Mice anesthetized with isoflurane and Beuthanasia-D (100 mg/kg, i.p.) were transcardially perfused with 20 mL ice-cold phosphate-buffered saline (PBS) followed by 50 mL of 4% paraformaldehyde (PFA) in PBS. Brains were extracted and post-fixed in 4% PFA for 24 hours at 4°C. On the next day, brains were transferred to 30% sucrose in PBS for overnight cryoprotection at 4°C.

#### Slicing

Coronal brain sections (40 μm thickness) were cut using a cryostat and stored at 4°C in a cryoprotectant solution (250 mL glycerol, 300 mL ethylene glycol, and 450 mL 1X PBS, with the pH adjusted to 6.7 using 1 M HCl).

#### Immunohistochemistry

Intrinsic GCaMP and ChRmine-oScarlet fluorescence was directly assessed without antibody staining. For the detection of NeuN and HA, an immunohistochemical protocol was employed: Sections were washed three times in PBST (PBS with 0.3% Triton X-100) for 10 min each. Sections were then incubated for 1 hr at room temperature in blocking buffer (PBS containing 0.3% Triton X-100 and 5% normal donkey serum) and subsequently incubated overnight at 4°C with the primary antibody diluted in blocking buffer. Primary antibodies used were rabbit anti-NeuN (Abcam REF# EPR12763, 1:200) and rat monoclonal anti-HA (clone 3F10; Roche, 0.1 µg/mL). Following primary antibody incubation, sections were washed three times in PBST for 30 minutes each. For secondary antibody incubation, sections were incubated for 2 hrs at room temperature with either donkey anti-rabbit IgG conjugated with Alexa-647 (Thermo Fisher Scientific Cat. No. A31573, used for NeuN detection) or donkey anti-rat IgG conjugated with Alexa-555 (Thermo Fisher Scientific Cat. No. A48270, used for HA detection).

Both were diluted 1:200 in blocking buffer. Finally, sections underwent four washes of 30 min each in PBST. Sections were mounted using ProLong Gold Antifade mounting medium (Thermo Fisher Scientific P10144).

#### Confocal imaging

Confocal images were acquired on a ZEISS LSM 880 scanning confocal microscope (Zen Black 2.3 software). Tile scans of entire brain slices were acquired using a 5X/0.2 NA objective lens (ZEISS) with a 2x digital zoom and stitched automatically within the Zen software. High-magnification images were obtained with a 20X/0.8 NA objective lens (ZEISS) and a digital zoom ranging from 0.7 to 1.5x. Optical sections were acquired with a pinhole set to 1 Airy Unit (AU). Laser power and detector gain were adjusted to maximize signal-to-noise ratio.

#### Quantification of confocal images

Images used to quantify co-labeled cells were acquired using a 20X/0.8NA objective lens and 1.5x digital zoom. FOVs in visual cortex spanning from L2/3 to upper L5, were chosen for quantification. Neuron segmentation was performed using Cellpose v2.2^78^ with the ‘cyto2’ model, followed by manual curation to correct segmentation errors. Co-labeling was determined using a custom script written in MATLAB (MathWorks, Inc.). A neuron was defined as co-labeled for any two markers (ChRmine-oScarlet, GCaMP, or NeuN) if the pixel overlap between their respective fluorescent channels exceeded a threshold of 30%.

## ACKNOWLEDGEMENTS

We thank Chris Roat for writing software for pre-processing of imaging data and Angie Gerbino, Alice Truong, Josh Moon and Vishal Mehta for technical assistance. The authors acknowledge support from the NIH BRAIN initiative (K.D.), the Gatsby Foundation (K.D.), and anonymous donors (K.D.), as well as a Human Frontiers in Science (HSFP) fellowship (A.D.) and a Swiss National Science Foundation (SNSF) fellowship (A.D.).

## AUTHOR CONTRIBUTIONS

A.D. and K.D. designed the study and experiments. A.D. performed all-optical experiments, with contributions from S.Q. and A.A. A.D., C.R., L.A.S., T.L.D., B.T., H.Z., and K.D. designed and developed transgenic reporter mice. A.D., A.A., and A.R. analyzed data, with input from S.Q. and S.G. A.R. and S.G. developed a computational model, with input from A.D. and A.A. C.R. performed cloning and generated viruses. B.E. performed surgeries, guided by A.D. and A.A. A.C. performed viral injections and histology, guided by A.D. Y.J. wrote software for online registration of two-photon experiments. A.D. and K.D. wrote the original draft of the manuscript. All authors reviewed and edited the manuscript. K.D. supervised all aspects of the work.

## MATERIAL AND DATA AVAILABITY

All materials, protocols, datasets and analysis code will be freely available to nonprofit institutions and investigators upon acceptance of the manuscript.

